# Multiple viral microRNAs regulate interferon release and signaling early during infection with Epstein-Barr virus

**DOI:** 10.1101/2020.12.03.393306

**Authors:** Mickaël Bouvet, Stefanie Voigt, Takanobu Tagawa, Manuel Albanese, Yen-Fu Adam Chen, Yan Chen, Devin N. Fachko, Dagmar Pich, Christine Göbel, Rebecca L. Skalsky, Wolfgang Hammerschmidt

**Affiliations:** Research Unit Gene Vectors, Helmholtz Zentrum München, German Research Center for Environmental Health and German Center for Infection Research (DZIF), Partner site Munich, Germany, Marchioninistr. 25, D-81377 Munich, Germany; Vaccine and Gene Therapy Institute, Oregon Health & Science University, 505 NW 185^th^ Ave, Beaverton, Oregon, USA; HIV and AIDS Malignancy Branch, Center for Cancer Research, National Cancer Institute, National Institutes of Health, Bethesda, Maryland, USA

## Abstract

Epstein-Barr virus (EBV), a human herpes virus, encodes 44 microRNAs (miRNAs), which regulate many genes with various functions in EBV-infected cells. Multiple target genes of the EBV miRNAs have been identified, some of which play important roles in adaptive antiviral immune responses. Using EBV mutant derivatives, we identified additional roles of viral miRNAs in governing versatile type I interferon (IFN) responses upon infection of human primary mature B cells. We also found that Epstein-Barr virus-encoded small RNAs (EBERs) and LF2, viral genes with previously reported functions in inducing or regulating IFN-I pathways, had negligible or even contrary effects on secreted IFN-α in our model. Data mining and Ago PAR-CLIP experiments uncovered more than a dozen of previously uncharacterized, direct cellular targets of EBV miRNA associated with type I IFN pathways. We also identified indirect targets of EBV miRNAs in B cells, such as TRL7 and TLR9, in the pre-latent phase of infection. The presence of epigenetically naïve, non-CpG methylated viral DNA was essential to induce IFN-α secretion during EBV infection in a TLR9-dependent manner. In a newly established fusion assay, we verified that EBV virions enter a subset of plasmacytoid dendritic cells (pDCs) and determined that these infected pDCs are the primary producers of IFN-α in EBV-infected peripheral blood mononuclear cells. Our findings document that many EBV-encoded miRNAs regulate type I IFN response in newly EBV infected primary human B cells in the pre-latent phase of infection and dampen the acute release of IFN-α in pDCs upon their encounter with EBV.

**Author summary:** Acute antiviral functions of all nucleated cells rely on type I interferon (IFN-I) pathways triggered upon viral infection. Host responses encompass the sensing of incoming viruses, the activation of specific transcription factors which induce transcription of IFN-I genes, the secretion of different IFN-I types and their recognition by the heterodimeric IFN-α/β receptor, the subsequent activation of JAK/STAT signaling pathways and, finally, the transcription of many IFN-stimulated genes (ISGs). In sum, these cellular functions establish a so-called antiviral state in infected and neighboring cells. To counteract these cellular defense mechanisms, viruses have evolved diverse strategies and encode gene products that target antiviral responses. Among such immune evasive factors are viral microRNAs (miRNAs) that can interfere with host gene expression. We discovered that multiple miRNAs encoded by Epstein-Barr virus (EBV) control over a dozen cellular genes that contribute to the anti-viral states of immune cells, specifically B cells and plasmacytoid dendritic cells (pDCs). We identified the viral DNA genome as the activator of IFN-α and question the role of abundant EBV EBERs, that, contrary to previous reports, do not have an apparent inducing function in the IFN-I pathway early after infection.

## Introduction

Epstein-Barr virus (EBV) is a DNA virus of the γ-herpesvirus family that establishes persistent, lifelong infection in the majority of the worldwide human population. Despite being mostly asymptomatic, EBV infection can cause infectious mononucleosis and lead to severe diseases such as Burkitt lymphoma (BL), diffuse large B cell lymphoma (DLBCL), nasopharyngeal carcinoma, and post-transplant lymphoproliferative disease (PTLD). PTLD and HIV-associated DLBCL develop in immunosupressed and immunocompromised individuals, highlighting the ability of the human immune system to keep EBV infection under control but the inability to reach total viral clearance.

EBV achieves latency in B lymphocytes through different phases which are characterized by the expression and downregulation of multiple viral gene products, ultimately decreasing the antigenic load of infected cells. EBV preferentially infects B lymphocytes and can even readily transform primary B lymphocytes into indefinitelly proliferating lymphoblastoid cell line (LCL) *in vitro*. Additional cell types in the blood, including plasmacytoid dendritic cells, have been reported to express viral gene products upon contact with EBV virions but cannot be transformed *in vitro* (Gujer et al., 2019 and references therein). Apart from viral proteins, the EBV genome encodes different classes of non-coding RNAs including two long non-coding RNAs, EBV-encoded small RNAs 1 and 2 (EBER1 and 2), circular RNAs (Ungerleider et al., 2018), and 44 mature microRNAs (miRNAs) distributed in clusters along the genome (Fig. 1).

**Figure 1.**
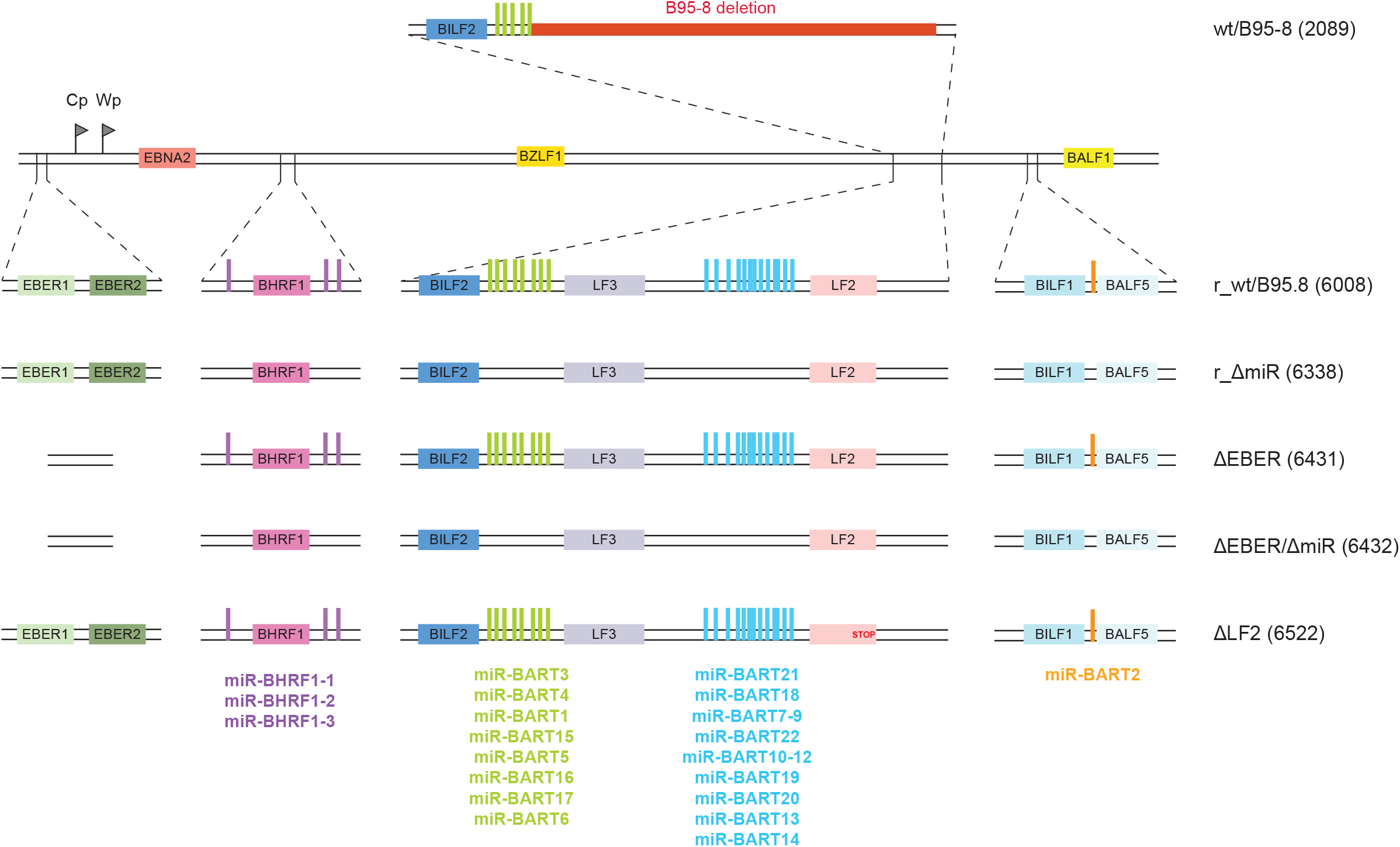
Construction of a reconstituted wild-type EBV genome and EBV mutants. A 11.8 kb fragment of the M-ABA strain complementing the deletion in EBV B95-8 was introduced by homologous recombination into wt/B95.8 (2089) to generate r_wt/B95.8 (6008) (for reconstituted wild-type). The 11.8 kb fragment contains the coding sequences of the LF2 and LF3 viral proteins and the majority of BART miRNAs, allowing for expression of these viral miRNAs at physiological levels. EBV r_wt/B95.8 (6008) was mutated to eliminate expression of either the viral miRNAs, the EBERs, or the LF2 protein. The r_ΔmiR (6338) strain was created by sequentially introducing three DNA fragments in which the viral pri-miRNA loci have been scrambled in order to prevent the formation of the characteristic hairpin structure recognized and processed by the ribonucleases Drosha and Dicer to generate mature miRNAs. The ΔEBER (6431) and ΔEBERΔmiR (6432) strains were created by insertional mutagenesis deleting the EBERs coding genes in r_wt/B95.8 and r_ΔmiR EBV derivatives, respectively. Finally, the ΔLF2 strain was created by introducing a stop codon in the r_wt/B95.8 (6008) strain preventing the translation of the LF2 protein. Supplementary Table 1 provides an overview of the different EBV strains.

In order to conduct a successful infection, viruses including EBV must evade or counteract the activation of type I interferons (IFN-I). IFN-I production and signaling is a two part system at the center of innate anti-viral immunity. Initially, specialized pattern recognition receptors (PRRs) detect pathogen-associated molecular patterns (PAMPs), which are distinct pathogen structures otherwise absent from non-infected, healthy cells. PRR engagement triggers activation of specific transcription factors which, in turn, induce transcription of IFN-I genes (13 IFN-α subtypes and IFN-β in humans), leading to production and secretion of type I IFNs. Secreted IFN-I binds to a single, heterodimeric IFN-α/β receptor on mammalian cells, thereby activating JAK/STAT signaling pathways and culminating with the transcription of many IFN-stimulated genes (ISGs). These gene products altogether establish a so-called antiviral state, thereby restricting the viral life cycle and/or orchestrating viral clearance in infected and neighbouring cells.

Due to their peculiar faculty to secrete massive amounts of IFN-I, plasmacytoid dendritic cells (pDCs) are a major factor in the IFN-I system. pDCs constitutively express major components of the IFN-I activation pathway like the TLR7 and TLR9 PRRs and the interferon regulatory factor 7 (IRF7) transcription factor. TLR7 and TLR9 are embedded in the endosomal membranes and respectively recognize single-stranded (ss)RNA or unmethylated CpG DNA in the endosomal lumen. Upon binding of PRRs, TLR7 and TLR9 signal through the MyD88 adaptor protein and activate the IRF7 transcription factor. There are hints that EBV is detected by pDCs, which respond to infection by secreting type I IFNs (Lim et al., 2007; Fiola et al., 2010; Quan et al., 2010; Severa et al., 2013). Whether the virus infects human pDCs and the mechanisms by which pDCs detect EBV are not fully understood.

It has been shown that the EBERs, which are abundant viral non-coding RNAs, can be recognized as PAMPs by the RIG-I, TLR3 and TLR7 PRRs and that pDCs further sense the presence of EBV through recognition of unmethylated viral genomic DNA by TLR9 (Fiola et al., 2010). Multiple intracellular sensors recognize exogenous cytosolic DNA, including DNA-dependent activator of IFN regulator factors (DAI) (Takaoka et al., 2007), DDX41 (Zhang et al., 2011), absent in melanoma 2 (AIM2) (Hornung et al., 2009), LSm14A (Li et al., 2012), IFN-gamma inducible factor 16 (IFI16) (Unterholzner et al., 2010), and cyclic GMP-AMP (cGAMP) synthase (cGAS) (Sun et al., 2013) which activates the adaptor protein STING (stimulator of IFN genes) to trigger IFN signaling. Human B cells express IFI16, cGAS, and downstream signaling components necessary to induce type I IFN; however, it has been demonstrated that B lymphocytes fail to recognize the presence of genomic EBV DNA through cGAS-STING (Gram et al., 2017).

EBV encodes several proteins that have been described as type I IFN antagonists and have attributed roles in counteracting anti-viral responses. In screening 150 EBV ORFs, EBV LF2, encoding a tegument protein, was identified as an inhibitor of IRF7 dimerization (Wu et al., 2009). BZLF1 also interferes with dimerization of IRF7 (Hahn et al., 2005) and blocks STAT1 tyrosine phosphorylation (Morrison et al., 2001) while BRLF1 controls both IRF3 and IRF7 (Bentz et al., 2010). The EBV kinase BGLF4 impedes IRF3 activity through direct interactions, consequently attenuating type I IFN signaling (Wang et al., 2009). Other viral proteins, such as BGLF5 (van Gent et al., 2011) inhibit production of TLR2 and TLR9.

In this study, we examined the relationship between EBV non-coding RNAs and activation of IFN response pathways. Here, we document that the EBV-encoded miRNAs contribute to the regulation of type I IFN response upon EBV infection, whereas EBERs and LF2 have a neglectable effect in infected human primary B cells. We identified several BART miRNAs which regulate genes involved in the IFN secretion and IFN response pathway. The presence of viral DNA was essential to induce the IFN-α secretion during EBV infection in a TLR9 dependent manner in PBMCs. In a newly established gp350:BlaM fusion assay, we verified that EBV virions enter a subset of pDCs and determined that these infected pDCs are the primary producers of type I IFNs in PBMCs.

## Results

### Cloning of a reconstituted wild-type EBV strain and mutant derivatives

To study the functions of EBV-encoded miRNAs, we constructed a recombinant EBV that expresses all viral miRNAs at their physiological levels and that can be genetically manipulated in bacteria to generate mutant derivatives. To do so, we used the wt/B95.8 (2089) recombinant virus (Fig. 1) available in our laboratory (Delecluse et al., 1998). This recombinant is based on the B95-8 strain of EBV (Miller and Lipman, 1973) into which a DNA fragment encoding a GFP gene (used for titration), a gene encoding resistance against hygromycin (used for selection in the EBV producer cells), a chloramphenicol acetyltransferase gene (used for selection in bacteria), and the mini-F factor replicon (enabling maintenance in bacteria) have been introduced (Delecluse et al., 1998). We further introduced a DNA fragment from the M-ABA field strain to restore viral genes that are deleted in EBV B95-8 (Bornkamm et al., 1980; Raab-Traub et al., 1980). This repaired B95-8 strain derivative, termed r_wt/B95.8 (6008) (Fig. 1) (Pich et al., 2019), encodes all known EBV miRNAs which are expressed from their genuine promoters at physiological levels. In addition, r_wt/B95.8 also carries the second copy of the lytic origin of DNA replication, *oriLyt*, (Hammerschmidt and Sugden, 1988) and expresses the LF1, LF2 and LF3 viral proteins from their coding sequences which are also absent in the genome of the B95-8 EBV strain (Supplementary Tab. 1).

Subsequently, we replaced the viral miRNA sequences in r_wt/B95.8 by scrambled sequences similar to our previous approach (Seto et al., 2010) preventing the expression of all viral miRNAs. This strain is called r_ ΔmiR (Fig. 1) (Pich et al., 2019).

The two viral non-coding EBER (<u>E</u>pstein–<u>B</u>arr virus-<u>e</u>ncoded small <u>R</u>NA) RNAs of 167 and 172 nucleotides in length have been implicated in the activation of type I IFN responses upon EBV infection in B cells (Samanta et al., 2006; Iwakiri et al., 2009). To evaluate their effects on type I IFN activation in our infection model, we deleted their coding sequences in r_wt/B95.8 and r_ΔmiR (Fig. 1). These two strains are called ΔEBER and ΔEBER/ΔmiR, respectively (Pich et al., 2019).

Finally, we mutated the gene encoding the LF2 viral protein that has been described as a type I IFN antagonist (Wu et al., 2009). We introduced a stop codon in the coding sequence of the LF2 gene to prevent its expression in r_wt/B95.8. This EBV strain is termed ΔLF2 (Supplementary Tab. 1).

To conclude, we present here a fully reconstituted EBV strain based on the B95-8 EBV genome. This recombinant EBV genome can be used to produce infectious virus, it can be conveniently genetically manipulated and presumably expresses all viral genes including all miRNAs at their physiological levels from their authentic promoters supported by their regulatory elements. We assume that, despite its chimeric composition, this new strain represents a more physiological model than the widely used B95-8 laboratory strain.

### Cytokines secretion by B lymphocytes infected by EBV mutants

We subsequently sought to determine the effects of the different mutations in the reconstituted r_wt/B95.8 EBV reference strain on the secretion of cytokines released from infected human B lymphocytes. Primary human B lymphocytes were infected *in vitro* with the five virus strains described above (r_wt/B95.8; r_ΔmiR; ΔEBER; ΔEBER/ΔmiR; ΔLF2). Upon infection with every virus strain tested, B cells grew in size and started to proliferate. Five days post-infection, infected cells were counted by flow cytometry and seeded at equal densities in fresh medium. Four days later, supernatants were collected and concentrations of IL-12, IL-6, IL-10, and IFN-α were measured by ELISA (Fig. 2).

**Figure 2.**
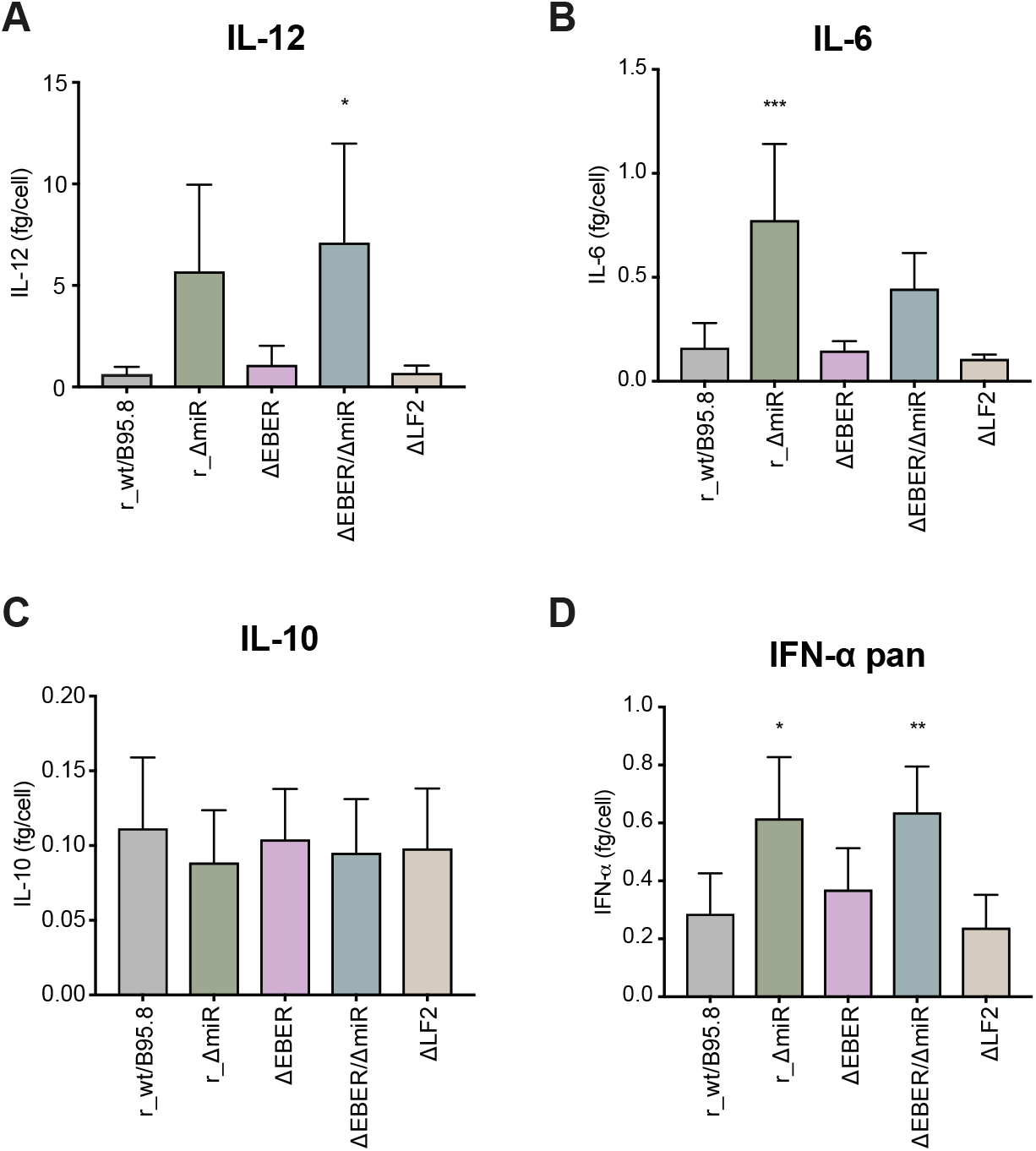
Cytokines secretion by infected primary B lymphocytes. Human primary B lymphocytes were infected with each of the five EBV strains at a multiplicity of infection (MOI) of 0.1 as described in Figure 1. Five days post-infection, EBV-infected cells were counted and seeded at the same density. Four days later, the cells were counted again, the culture supernatants were collected, and cytokine levels were analyzed by ELISA. The supernatant concentrations of IL-12 **(A)**, IL-6 **(B)**, IL-10 **(C)** and IFN-α pan **(D)** are normalized to the final cell number and reported as fg/cell.

We showed previously that EBV-encoded miRNAs regulate the expression of human IL-12 (Tagawa et al., 2016). As shown in Figure 2A, B lymphocytes infected with r_ΔmiR secreted more IL-12 than cells infected with r_wt/B95.8, consistent with prior experiments using the B95-8 strain wt/B95.8 (2089) (Fig. 1) (Tagawa et al., 2016). The deletion of the EBERs or LF2 did not influence IL-12 secretion. Moreover, when the cells were infected with the double mutant ΔEBER/ΔmiR, IL-12 was secreted at concentrations similar to r_ΔmiR infected B cells. These data confirm that the EBV-encoded miRNAs are responsible for altered IL-12 levels in newly infected B lymphocytes while the EBERs and LF2 do not affect IL-12 production.

We have further documented that EBV-encoded miRNAs inhibit the secretion of IL-6 but not IL-10 in the B95-8 context (Tagawa et al., 2016). As shown in Figure 2B, the absence of the viral miRNAs in cells infected with r_ΔmiR led to a substantial increase in IL-6 secretion when compared to r_wt/B95.8-infected cells. This finding demonstrates the importance of the viral miRNAs to regulate IL-6 secretion upon EBV infection. Interestingly, it has been suggested that EBER2 also plays a role in IL-6 secretion (Wu et al., 2007). We did not observe any difference in IL-6 secretion by cells infected with r_wt/B95.8 or ΔEBER, but the ΔEBER/ΔmiR-infected cells did secrete less IL-6 than the r_ΔmiR-infected cells (Fig. 2B). Results from our infection model suggest that EBERs may stimulate the expression of IL-6 to some extent, but the viral miRNAs largely repress IL-6 secretion. As for IL-12 secretion, ΔLF2 behaved very similar to r_wt/B95.8 (Fig. 2B).

Concerning IL-10 secretion, none of the mutations clearly altered the cytokine secretion by infected B lymphocytes (Fig. 2C). It has to be noted that our ELISA quantification cannot differentiate between human and viral IL-10 encoded by EBV (Moore et al., 1990). Thus, we cannot rule out that human IL-10 expression is regulated by viral miRNAs, EBERs, or LF2 but is compensated by the expression of viral IL-10 encoded by the viral BCRF1 gene.

We next asked whether mutations in the EBV mutant derivatives would impact the secretion of IFN-α by EBV-infected B lymphocytes. As shown in Figure 2D, the absence of viral miRNAs led to increased secretion of IFN-α by r_ΔmiR-infected cells compared to r_wt/B95.8-infected cells. Surprisingly, we did not observe any decrease in secreted IFN-α when comparing ΔEBER-to r_wt/B95.8-infected cells and ΔEBER/ΔmiR-to ΔmiR-infected cells. These data are puzzling since EBERs have been reported to interact with the PRR RIG-I and act as type I IFN inducers (Samanta et al., 2006; Samanta et al., 2008; Iwakiri et al., 2009; Iwakiri, 2014; Duan et al., 2015). Additionally, ΔLF2-infected cells did not show any significant difference compared to r_wt/B95.8-infected cells despite the fact that LF2 has been shown to interact with IRF7 and block the expression of type I IFN (Wu et al., 2009). These discrepancies could be explained by differences in the experimental conditions. What we can ascertain is that, in our experiments with primary cells, the EBERs are not necessary for the expression and secretion of IFN-α by EBV-infected B lymphocytes. Moreover, independent of the presence or absence of the EBERs, the viral miRNAs significantly impact the secretion of IFN-α.

Taken together our data indicate that EBV-encoded miRNAs prevent the secretion of IL-12, IL-6 and IFN-α in newly infected primary human B cells. Whereas the EBERs seem to be partially responsible for the activation of IL-6, their deletion did not cause any significant difference in the secretion of IL-12, IL-10 or IFN-α. We also failed to observe a contribution of LF2 to the secretion of the cytokines analyzed.

### Identification of cellular transcripts regulated by EBV-encoded miRNAs

Based upon the phenotypes we observed above, we hypothesized that EBV-encoded miRNAs regulate the expression of host cellular genes involved in multiple aspects of the IFN pathways, such as PAMP sensing, production of type I IFN, and/or induction of secondary cytokine signals that encompass innate anti-viral responses. To initially investigate targets of EBV miRNAs that may be involved in these pathways, we first used *in silico* prediction tools (Garcia et al., 2011) to scan human 3’UTRs for the presence of canonical 7mer and 8mer miRNA seed-match sites. Candidates were cross-referenced with the literature to determine putative targets associated with PRR activation and type I IFN signaling. Among these, we identified potential binding sites for EBV miRNAs in the 3’UTRs of *DDX58/RIG-I, RSAD2/Viperin, OAS2*, and *FYN*.

To biochemically define EBV miRNA binding sites in human protein-coding transcripts, we performed PAR-CLIP experiments on latently infected DLBCL cell lines (IBL1, IBL4, and BCKN1) which express the full spectrum of EBV BHRF1 and BART miRNAs. Cells were cultured in the presence of 4-thiouridine to label RNAs as previously described (Skalsky et al., 2012). RISC-associated RNAs were immunopurified with antibodies to Argonaute (Ago) proteins and subjected to high-throughput sequencing. Following alignment to a human reference genome, reads were analyzed using two CLIP-seq pipelines (PARalyzer and PIPE-CLIP) to define high resolution Ago interaction sites (Corcoran et al., 2011; Chen et al., 2014). To identify host mRNAs reproducibly targeted by EBV miRNAs, we expanded our analysis to include previously published Ago PAR-CLIP datasets from EBV B95-8 and wildtype LCLs and EBV/KSHV+ PEL (Supplementary Tab. 2, Supplementary Fig. 1) (Al Tabaa et al., 2009; Gottwein et al., 2011; Skalsky et al., 2012; Majoros et al., 2013; Skalsky et al., 2014). Derived Ago interaction sites mapping to 3’UTRs of protein coding transcripts were scanned for canonical EBV miRNA seed matches (>=7mer1A). In total, we identified 4,010 individual genes regulated at the 3’UTR level by EBV miRNAs, with 340 genes common to all EBV-infected B cell types investigated (DLBCLs, LCLs, PEL) (Supplementary Fig. 1E).

To focus on EBV miRNA targets functionally relevant to type I IFN pathways, we then interrogated Ago PAR-CLIP data using Ingenuity Pathway Analysis (IPA, Qiagen). 3,976 of the 4,010 3’UTR target genes were mapped by IPA. Enriched signaling pathways included ‘JAK/Stat Signaling’ and ‘Role of PKR in Interferon Induction and Antiviral Response’ (Supplementary Fig. 1F). Among the target genes related to IFN pathways, we identified *JAK1, IRF9, STAT1, IRAK2, IKBKB*, and interferon receptors (*IFNAR1* and *IFNAR2*). Together, these data reinforce the observation that EBV miRNAs play a major role in regulating innate immune response pathways.

### EBV-encoded miRNAs regulate genes involved in the induction of type I IFN

To confirm miRNA interactions, we first selected targets with relevance to induction of type I IFN (*DDX58/RIG-I, RSAD2/Viperin, OAS2, FYN*, and *IRAK2*) and cloned the 3’UTRs of these genes into reporter plasmids. Reporter plasmids were co-transfected into 293T cells with or without an EBV miRNA-expressing plasmid and luciferase expression was assessed (Fig. 3). When we observed miRNA mediated inhibition of luciferase expression, we mutated the predicted seed-sequences in the reporter plasmid and verified that the miRNA expression did not have an effect on the expression of luciferase (Supplementary Fig. 2).

**Figure 3.**
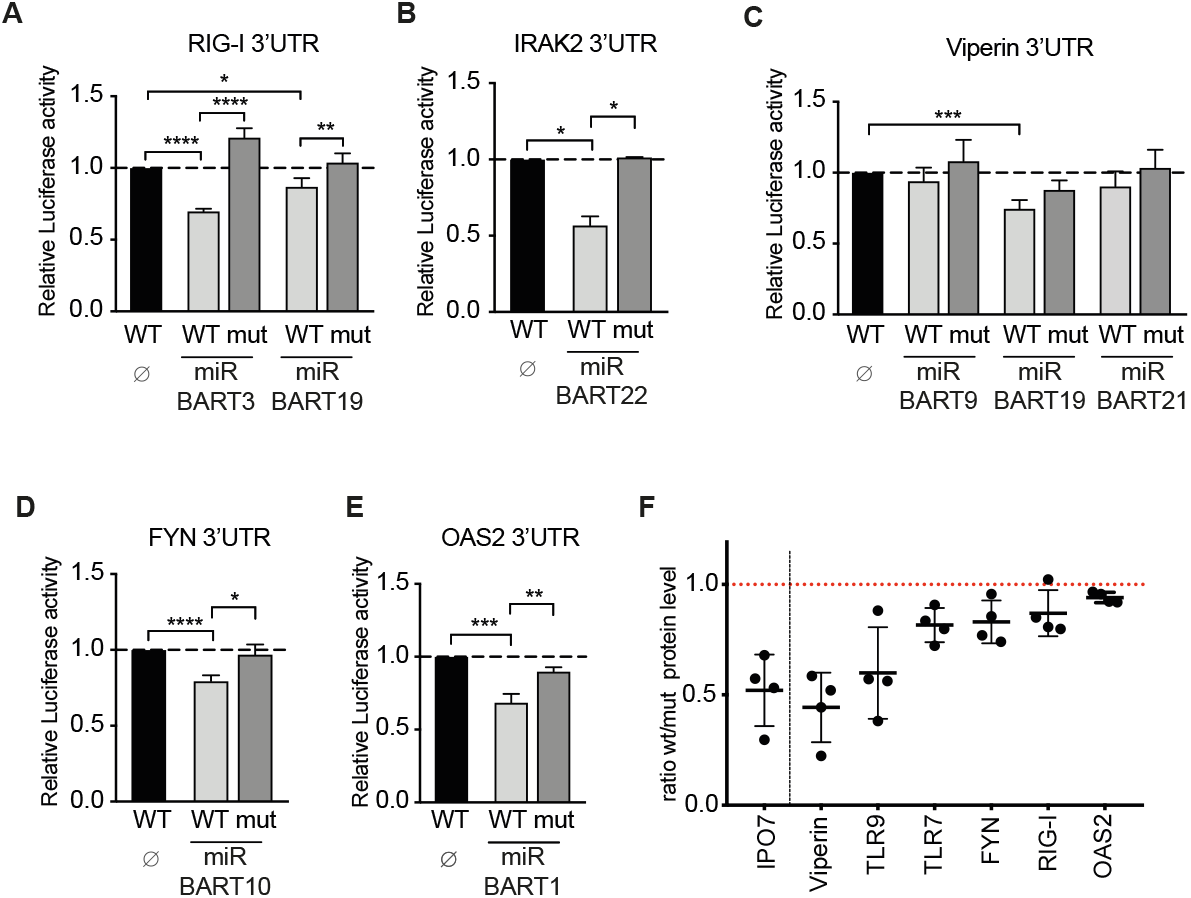
EBV-encoded miRNAs regulate the expression of cellular genes and proteins involved in type I IFN activation. **(A-E)** 293T cells were co-transfected with the indicated wild-type luciferase reporter plasmids (WT) for RIG-I/DDX58 (A), IRAK2 (B), Viperin/RSAD2 (C), FYN (D) and OAS2 (E) or reporter plasmid in which the predicted seed sequence was mutated (mut) together with or without an miRNA-encoding plasmid. The luciferase expression in these cells was assessed and normalized to lysates from cells co-transfected with the wild-type 3’-UTR reporter and empty miRNA expression plasmid (∅). P-values were calculated using the one-way ANOVA test. *, P < 0.05; **, P < 0.01; ***, P < 0.001; ****, P < 0.0001. **(F)** Human primary B lymphocytes were infected with wild-type EBV (r_wt/B95.8) or EBV devoid of its miRNAs (r_ ΔmiR). Five days post-infection, EBV-infected cells were counted and seeded at the same density. Four days later, the cells were lysed and the lysates were subjected to quantitative Western Blot analysis using the Western Blot Stain-free TGX Biorad Normalization approach (Bio-Rad). Blots were probed with antibodies against IPO7, Viperin/RSAD2, TLR9, TLR7, FYN, RIG-I/DDX58 and OAS2. Protein levels were quantified and used to normalize the levels of the specific protein signals. Ratios of protein levels in cells infected with wildtype virus versus cells infected with mutant virus are shown. Reported are the results of four independent biological replicates.

We initially tested miR-BART3 and miR-BART19 as regulators of the RIG-I 3’UTR (Fig. 3A). It has recently been reported that another EBV miRNA, miR-BART6, acts as a regulator of RIG-I expression and prevents the expression of type I IFN (Lu et al., 2017). Our findings confirm that EBV-encoded miRNAs control the expression of the RIG-I PRR. As the virus has evolved mechanisms to attenuate RIG-I expression, this suggests that RIG-I is involved in efficiently detecting EBV infection. Interestingly, even though the EBERs have so far been described as the primary virus moiety activating RIG-I, our data suggest that there are viral PAMPs recognized by the RIG-I helicase other than EBERs since the ΔEBER-infected cells still secrete IFN-α at levels similar to its wildtype predecessor r_wt/B95.8 (Fig. 2D).

As shown in panels B to E of Fig. 3, we found EBV-encoded miRNAs to also regulate the 3’UTRs of *IRAK2, RSAD2/Viperin, FYN*, and *OAS2*. These genes have been described as important factors for the production of type I IFN as well as activation of human pDCs. IFN-inducible *RSAD2/Viperin* promotes IFN-α secretion upon TLR7 and TLR9 activation (Saitoh et al., 2011), while *IRAK2* is essential for the expression of type I IFN initiated by TLR7 in pDCs (Kawagoe et al., 2008; Flannery et al., 2011; Wang et al., 2015). A study also showed the importance of a functional *IRAK2* for the late stage of cytokines production including IFN-α in murine pDCs (Pauls et al., 2013). Modulating the expression of *IRAK2* by EBV miRNAs could thus contribute to regulating the release of IFN-α by pDCs via the inhibition of the TLR7 pathway.

*OAS2* (2’-5’-oligoadenylate synthetase 2) is an ISG that specifically recognizes dsRNA structures and synthesizes 2’-5’-oligoadenylate structures that activate RNAseL. RNaseL subsequently degrades cellular and viral RNAs and thus inhibits translation in the infected cell. We show that miR-BART1 can regulate the 3’UTR of OAS2 in Figure 3E; however, compared to RSAD2/Viperin, TLR9, TLR7, RIG-I and FYN protein levels which inversely correlated with EBV miRNA presence, OAS2 protein levels were barely upregulated in r_ΔmiR-infected cells (Fig. 3F). Together, these data show that EBV-encoded miRNAs inhibit expression of multiple cellular factors involved in activation, expression, and secretion of type I IFN.

### EBV-encoded miRNAs regulate the response to type I IFN signaling

Upon secretion, type I IFN interacts with the heterodimeric IFN-α/β receptor (IFNAR) and initiates a signaling cascade through the JAK-STAT pathway. To determine if EBV-encoded miRNAs impact type I IFN responses in addition to inhibiting IFN production, we used a reporter luciferase system whereby type I IFN was added exogenously in the presence of individual EBV miRNAs. We constructed a luciferase reporter plasmid p6898 with the improved firefly luciferase (*luc2*) gene under control of a chimeric promoter comprising part of the ISG Mx2 promoter and a repetition of five canonical ISREs (Interferon-Stimulated Response Element). A renilla luciferase gene under the control of the weak TK promoter was used as an internal transfection control for subsequent data normalization. We concomitantly transfected this vector into 293T cells with individual expression vectors for EBV-encoding miRNAs or an empty vector as a control. 24 h post-transfection, cells were treated with type I IFN and luciferase signals were quantified one day later. As a positive control, we used a miRNA expression vectors for human miR-373 (Fig. 4A), which regulates the expression of *JAK1* and *IRF9* (Mukherjee et al., 2015), and is thus expected to reduce the response to exogenous IFN. We also cloned and tested the IE1 gene from human cytomegalovirus (HCMV). IE1 is a very potent inhibitor of type I IFN signaling (Piganis et al., 2011) and indeed efficiently suppressed luciferase expression in our assay (Fig. 4A).

**Figure 4.**
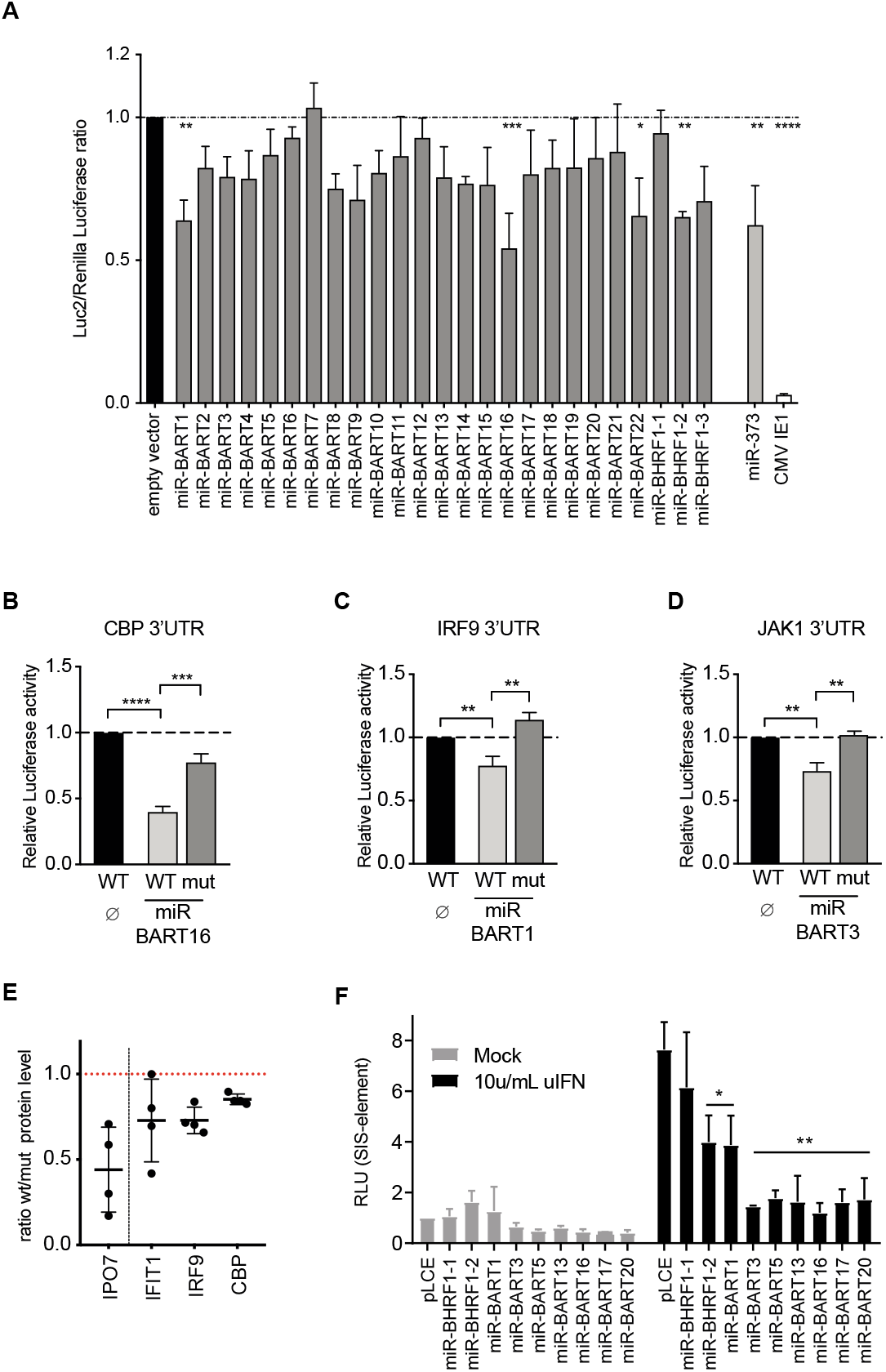
Interferon-stimulated genes (ISG) are regulated by EBV-encoded miRNAs. **(A)** 293T cells were co-transfected with individual EBV miRNA expression vectors or an empty vector as control and a reporter plasmid (p6898) containing a luc2 gene under control of a chimeric promoter consisting of the ISG Mx2 promotor and five canonical ISREs. After 24 h, cells were treated with type I IFN and luciferase signals were measured on the next day. As control, an expression vector encoding human miRNAs (miR-373) was used (p6481). A further control was the IE1 gene from CMV, which is known to inhibit type I IFN signaling (Piganis et al., 2011). P-values were calculated by one-way ANOVA test. *, P < 0.05; **, P < 0.01; ***, P < 0.001; ****, P < 0.0001. **(B-D)** 293T cells were co-transfected with the indicated wild-type luciferase reporter plasmids (WT) for CBP, IRF9, and JAK1 or reporter plasmids in which the predicted seed sequences were mutated (mut) together with or without a miRNA-encoding plasmid. The luciferase expression in these cells was assessed and normalized to lysates from cells co-transfected with the wild-type 3’-UTR reporter and empty plasmid (∅). P-values were calculated using the one-way ANOVA test. *, P < 0.05; **, P < 0.01; ***, P < 0.001; ****, P < 0.0001. **(E)** Human primary B lymphocytes were infected with wild-type EBV (r_wt/B95.8) or EBV devoid of its miRNAs (r_ ΔmiR). Five days post-infection, the transformed EBV-infected cells were counted and seeded at the same density. Four days later, the cells were lysed and the lysates were subjected to quantitative Western Blot analysis using the Western Blot Stain-free TGX Biorad Normalization approach (Bio-Rad) and antibodies against IPO7, IFIT1, IRF9 and CBP. The applied protein amounts were analyzed and used to normalize the levels of the specific protein signals. The ratios of protein levels of cells infected with wildtype virus versus cells infected with mutant virus are shown. The figure summarizes the results of four independent replicates. (**F)** HEK293T cells were co-transfected with a pLCE based EBV miRNA expression vector, a SIS-inducible element (SIE) responsive firefly luciferase reporter and a pRL-SV40 renilla luciferase reporter as internal control. 48 hrs after transfection, cells were either mock treated or treated with 10 unit per mL universal type I interferon (uIFN) for 6 hrs, and analyzed for luciferase activity. Values are normalized to mock treated cells and reported relative to pLCE control. Shown are the averages of three to four independent experiments performed in triplicate. Data were analyzed using Student’s t-test, *p<0.05, **p<0.02.

Through this functional screen, we identified four EBV-encoded miRNAs that significantly attenuated the response to exogenous type I IFN (miR-BART1, miR-BART16, miR-BART22 and miR-BHRF1-2) as well as multiple other EBV miRNAs that moderately attenuated responses (Fig. 4A). miR-BART16 has recently been shown to regulate the expression of CREB-binding protein (*CBP*) and thereby inhibits the induction of ISGs (Hooykaas et al., 2017). Targeting of the *CBP* 3’UTR was confirmed in dual-luciferase 3’UTR reporter assays (Fig. 4B).

To investigate cellular targets that are likely responsible for the phenotypes observed for the other EBV miRNAs, we examined miRNA targetome datasets (Supplementary Fig. 1). Notable targets included *JAK1* and *IRF9* which are directly involved in JAK/STAT signaling in response to type I IFN. IRF9 is part of the ISGF3 (Interferon-stimulated gene factor 3) complex together with phosphorylated STAT1 and STAT2 and is required to drive the expression of ISGs upon type I IFN signaling (Qureshi et al., 1995). Limiting the expression of IRF9 could be a strategy employed by EBV to prevent ISGF3 complex association and the transcription of ISGs. JAK1 is a kinase and its association with the type I IFN receptor is required to transduce the signal upon ligation of IFN on the receptor. We cloned 3’UTRs of these and other genes into dual-luciferase reporter plasmids and tested them against individual EBV miRNAs. Notably, miR-BART1 inhibited luciferase expression from the *IRF9* reporter (Fig. 4C) while miR-BART3 reduced activity of the *JAK1* reporter (Fig. 4D). miR-BART3 had a moderate effect on luciferase expression in our IFN-response assay (Fig. 4A) that could be explained by this finding. We furthermore found that miR-BART2 regulates the *JAK2* 3’UTR, miR-BART17 regulates the *IKKB* 3’UTR, while miR-BART3 and miR-BART22 regulate *MAP3K2* (Supplementary Fig. 3A-C). In our dual-luciferase reporter assays, we could further observe a suppressive effect of EBV-encoded miRNAs on reporter plasmids for *IFNAR1, IRF9, CHUK, ZCCHC3, VAV2, RAC1* and *VAMP3*, which are genes with well-known functions in type I interferon signaling (Supplementary Fig. 3E-K).

To confirm that EBV miRNAs indeed attenuate IFN-mediated activation of the JAK-STAT pathway, we tested a STAT-responsive luciferase reporter harboring SIS-inducible elements (SIE) in the presence of EBV miRNAs in 293T cells (Fig. 4F). Consistent with inhibition of JAK1 and IRF9, both miR-BART1 and miR-BART3 suppressed SIE reporter activity following treatment with IFN. qRT-PCR analysis of IFN-treated 293T cells further revealed that ISG54 and ISG56 transcript levels were reduced in the presence of miR-BART1 and miR-BART3 (Supplementary Fig. 4). These data demonstrate that EBV-encoded miRNAs inhibit the expression of cellular genes involved at every step of the type I IFN pathway, including the response to type I IFN and induction of ISGs.

### EBV-encoded miRNAs regulate the expression of proteins involved in the secretion of type I IFNs

From the identified targets of EBV-encoded viral miRNAs, we validated some at the level of protein expression. Primary human B cells were infected with the r_wt/B95.8 or r_ ΔmiR EBV strains for 9 days. The cells were lysed and the protein-adjusted cell lysates were analyzed for the expression of the indicated proteins by Western Blot immunodetection. As a positive control, we monitored the expression of IPO7 (Fig. 3F and 4E), which is a published target of EBV miRNAs (Dölken et al., 2010). RSAD2/Viperin, TLR9, TLR7, FYN, RIG-I/DDX58 and OAS2, which are known to be involved in the type I IFN induction, are downregulated in r_wt/B95.8 infected B cells (Fig. 3F) probably directly or indirectly by EBV-encoded miRNAs. Furthermore IFIT1, IRF9 and CBP, which are involved in the type I IFN response, are affected by EBV-encoded miRNAs (Fig. 4E).

### IFN-α secretion by PBMCs and isolated pDCs infected with mutant EBV strains

We next asked whether PBMCs infected with the designed EBV mutants secrete IFN-α. We infected human PBMCs with the five EBV strains (Supplementary Tab. 1) at three different multiplicities of infection (MOI). Culture supernatants were collected 20 h post-infection and secreted IFN-α was quantified by ELISA. As shown in Figure 5A, PBMCs released IFN-α in a dose-dependent manner upon EBV infection. At the lowest MOI, r_wt/B95.8-, ΔEBER- and ΔLF2-infected cells released a similar IFN-α concentration while the absence of miRNAs in r_ ΔmiR- and ΔEBER/ΔmiR-infected cells led to higher IFN-α secretion. At higher MOIs (0.05 and/or 0.1), the differences between the individual EBV mutants became more marked. Under these conditions, ΔEBER- and ΔLF2-infected cells released slightly more IFN-α than r_wt/B95.8-infected cells (Fig. 5A) suggesting that non-coding EBERs and the gene product of LF2 could dampen expression and/or secretion of IFN-α. Additionally, ΔEBER/ΔmiR-infected cells released more IFN-α than r_ ΔmiR-infected cells, again indicating that EBERs might be involved in controlling IFN-α secretion.

**Figure 5.**
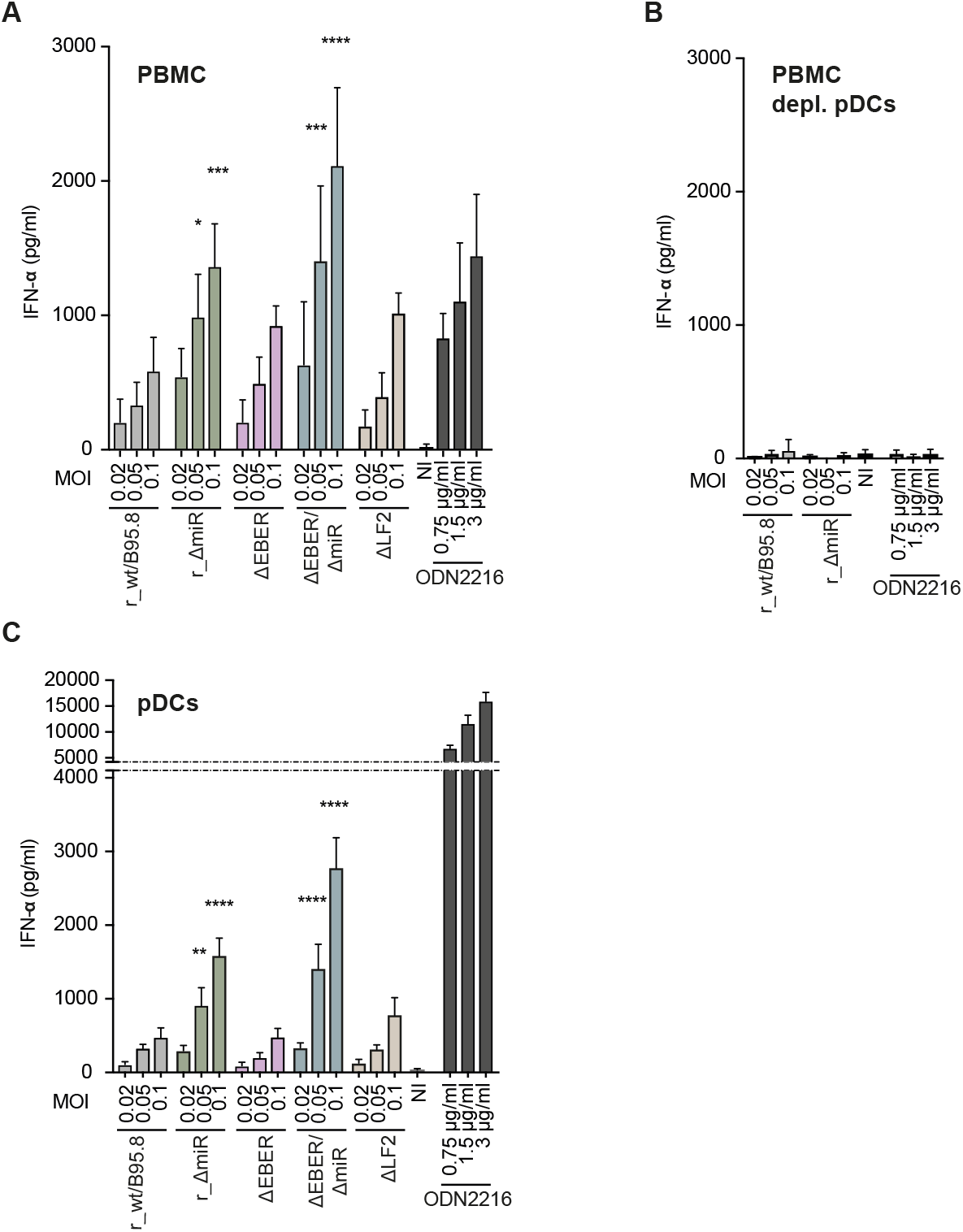
Plasmacytoid dendritic cells (pDCs) sense EBV infection and respond by secreting IFN-α. **(A)** PBMCs were infected with the five EBV strains described in Figure 1 and listed in Supplementary Table 1. The culture supernatants were collected 20 h post-infection and IFN-α concentrations were quantified by ELISA. P-values were calculated using the one-way ANOVA test. *, P < 0.05; ***, P < 0.001; ****, P < 0.0001. **(B)** pDCs were depleted from PBMCs using anti-CD304-coupled microbeads. The pDC-depleted PBMCs were infected with r_wt/B95.8 and r_ΔmiR using the same conditions as in panel A and IFN-α concentrations were determined by ELISA. **(C)** pDCs were isolated from buffy coats by negative selection and infected with the five EBV strains described in Figure 1. Cell culture supernatants were collected 20 h post-infection and IFN-α concentrations were determined by ELISA. P-values were calculated using the one-way ANOVA test. **, P < 0.01; ****, P < 0.0001. Reported are the results of four independent biological replicates.

It has been reported that plasmacytoid dendritic cells (pDCs) detect EBV infection and respond by releasing type I IFN (Lim et al., 2007; Fiola et al., 2010; Quan et al., 2010; Severa et al., 2013). In an attempt to assess the role of viral miRNAs on the sensing of infectious EBV by pDCs, we first depleted pDCs from PBMCs. We infected pDC-depleted PBMCs with r_wt/B95.8 and r_ ΔmiR using the same conditions as above and collected the culture supernatants 20 h post-infection. IFN-α release was quantified by ELISA. As shown in Figure 5B, the depletion of pDCs resulted in an almost complete loss of IFN-α secretion by the remaining cells, including B lymphocytes. This finding confirms data by Severa and colleagues (Severa et al., 2013) and supports the idea that pDCs are primarily responsible for the immediate IFN-α release when PBMCs are infected with EBV (Fig. 5A). B lymphocytes barely released IFN-α immediately after PBMC infection (Fig. 5B). Moreover, in the pre-latent phase, EBV-infected B lymphocytes secreted IFN-α, but at a much lower concentration (Fig. 2D) compared to pDCs (Fig. 5C).

We then isolated pDCs and infected the cells with the five EBV strains. As shown in Figure 5C, r_ ΔmiR-infected pDCs secreted more IFN-α than r_wt/B95.8-infected pDCs, consistent with our findings in PBMCs. ΔEBER- and ΔLF2-infected pDCs released IFN-α similar to r_wt/B95.8-infected cells whereas ΔEBER/ΔmiR-infected pDCs again released the highest amount of IFN-α suggesting that EBERs and viral miRNAs might even work collaboratively and reduce activation of the type I IFN pathway. The absence of LF2 did not seem to cause a marked effect on the secretion of IFN-α by pDCs.

Thus far, our experiments suggest that EBV-encoded miRNAs are directly involved in the regulation of type I IFN responses, especially in regulating gene expression of different pathway components. The overall impact of EBV miRNAs on direct IFN-α release by pDCs appears to be moderate, suggesting that EBV miRNAs act to reduce amplification of the IFN response in the context of PBMCs.

### IFN-α release by EBV infected cells is triggered by viral DNA and depends on TLR9 signaling

We identified pDCs as the main producers of IFN-α upon EBV infection, which is in line with published literature (Quan et al., 2010; Severa et al., 2013; Gujer et al., 2019). As described previously, pDCs express TLR7 and TLR9 which sense viral RNA and DNA, respectively, for secreting type I IFNs (Colonna et al., 2004; Reizis, 2019). These TLRs are found in the endosomal compartment and seem to be shifting between endosomes and lysosomes (Ahmad-Nejad et al., 2002). We wanted to analyze if viral DNA is necessary and sufficient for the secretion of IFN-α by PBMCs upon EBV infection. EBV DNA is very rich in CpG dinucleotides, which are non-methylated in virions (Fig. S3 in Kalla et al., 2010). Similar to bacterial DNA, unmethylated virion DNA is an ideal pattern for recognition by TLR9.

Towards this end, we infected PBMCs with the r_ ΔmiR EBV strain or incubated the cells with adjusted concentrations of virus-like particles (VLPs), which are free of viral DNA (Hettich et al., 2006). 20 h post infection supernatants were collected and levels of IFN-α were measured. As expected, infection with r_ ΔmiR led to a dose-dependent secretion of type I IFN, but PBMCs treated with VLPs did not secrete detectable levels of IFN-α (Fig. 6A, Supplementary Fig. 5). To confirm that the presentation and sensing of viral DNA by pDCs is important for type I IFN release after EBV infection, we treated EBV-infected PBMCs with chloroquine. Chloroquine is an inhibitor of lysosomal functions and blocks the activation of endosomal TLRs including TLR7 and TLR9. Chloroquine treatment of EBV infected PBMCs led to a complete block of IFN-α release compared with PBMCs infected with EBV in the absence of the lysosome inhibitor (Fig. 6B).

The synthetic oligo deoxyribonucleotide ODN2088 is an antagonist, which inhibits the TLR7/8/9 mediated responses in cells. The corresponding oligonucleotide ODN2087 is a TLR7/8 antagonist, but also acts as a TLR9 antagonist control, as it has no effect on TLR9 signaling. To find out whether the IFN secretion in EBV-infected PBMCs is dependent on TLR9 as a sensor for unmethylated CpG-DNA, we infected human PBMCs with EBV r_ ΔmiR in the presence of the TLR9 inhibitor ODN2088 or its ODN2087 control oligonucleotide. The inhibition of TLR9 nearly completely blocked the IFN release of EBV-infected PBMCs (Fig. 6C). When the EBV-infected PBMCs were treated with ODN2087 control, a slight drop in IFN-α levels compared to the untreated cells was noticed (Fig. 6C).

**Figure 6.**
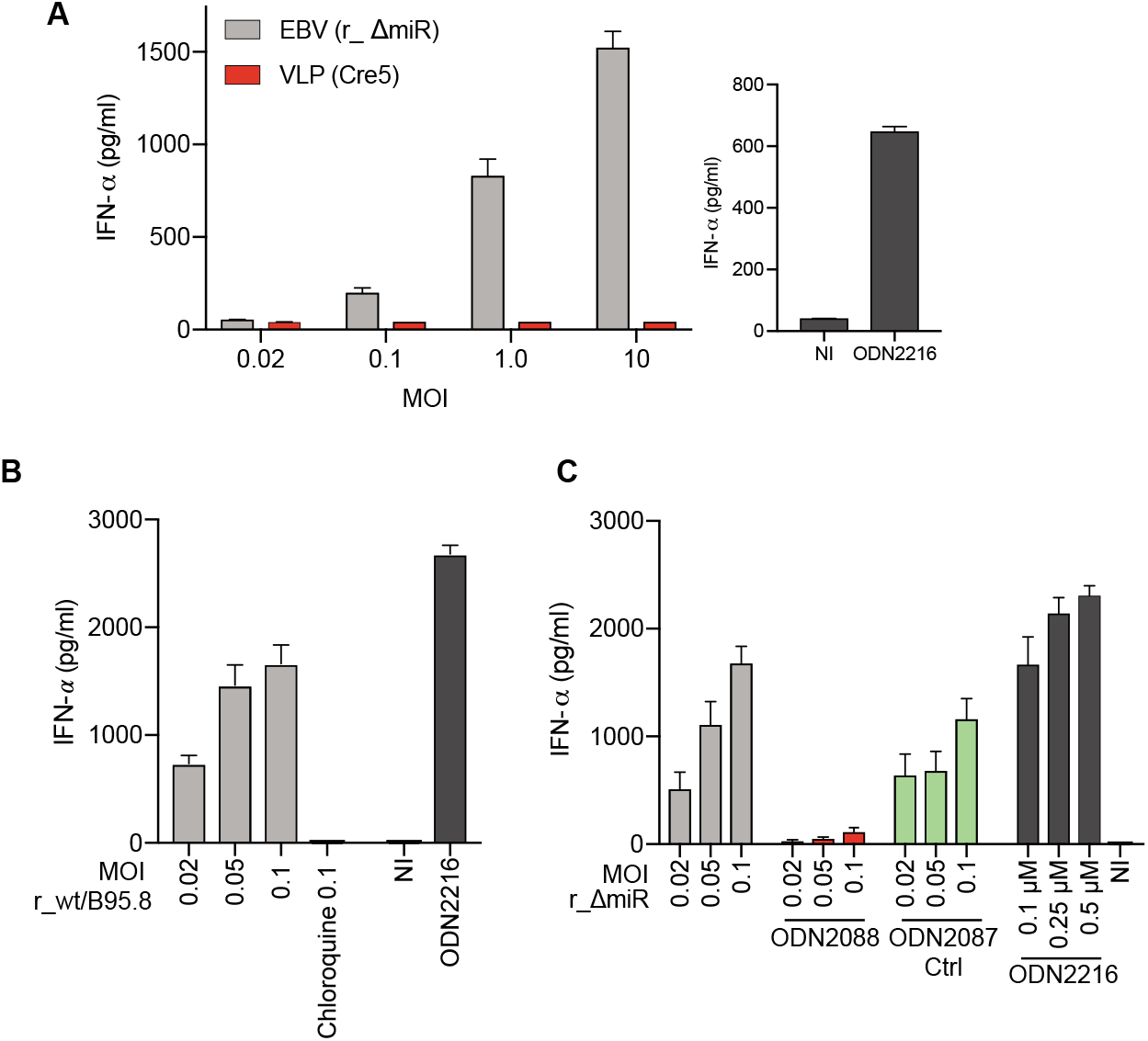
EBV-induced IFN-α release by PBMCs depends on viral DNA and TLR9 signaling. **(A)** PBMCs were infected with r_ΔmiR EBV or with virus-like particles (VLP Cre5), which do not contain viral DNA. Non-infected (NI) cells and cells treated with the TRL9 agonist ODN2216 served as negative and positive controls, respectively. The culture supernatants were collected 20 h post-infection and the IFN-α levels were quantitated by ELISA. Mean and standard deviation of three to four biological replicates are shown. **(B)** PBMCs were infected with EBV r_wt/B95.8 with different MOIs as indicated with or without treatment with 100 μM chloroquine. The culture supernatants were collected 20 h post-infection and the IFN-α levels were quantitated by ELISA. **(C)** PBMCs were infected with EBV r_ΔmiR using different MOIs as indicated. The cells were left untreated or they were treated with 0.25 μM TLR9 antagonist ODN2088 or 0.25 μM ODN 2088 control oligonucleotide, termed ODN 2087. ODN2087 acts as a TLR9 antagonist control. Cells treated with different concentrations of the TRL9 agonist ODN2216 served as positive control, non-infected (NI) cells act as background control. The culture supernatants were collected 20h post-infection and the IFN-α levels were quantitated by ELISA. Reported are the results of four independent biological replicates.

Together, these results demonstrate that IFN-α release during EBV infection of PBMCs depends on viral DNA and its detection by TLR9.

### EBV-infected pDCs are the main producers of IFN-α

As shown above, type I IFN released immediately after EBV infection of PBMCs almost exclusively originates from pDCs. Our data shown in Figure 5 suggest that viral miRNAs and probably also the two non-coding EBERs modulate the release of IFN-α. Both classes of RNAs are contained in the tegument (i.e. the space between envelope and capsid) in EBV virions and can be delivered to EBV target cells (Pegtel et al., 2010; Jochum et al., 2012; Baglio et al., 2016). IFN-α release by pDCs implies that these cells endocytose EBV particles as B cells do. For pDCs to sense virion DNA and present it to the TLR9 PPR, it is mandatory that EBV particles release their virion DNA probably when endosomes reach the degradative lysosome. In this scenario, it is unclear how viral miRNAs contained in the tegument might modulate the type I IFN response unless they can be delivered into the cytoplasm of pDCs (or any other cell type where they act in conjunction with the RNA-induced silencing complex) prior to lysosomal degradation. We therefore asked if EBV can infect pDCs as has been claimed previously (Severa et al., 2013) and developed an assay to determine the cell types with which EBV particles fuse to deliver their cargo into the cytoplasm.

We turned to wt/B95.8 (2089) producer cells (Delecluse et al., 1998) (Fig. 1) as they release very high amounts of infectious EBV upon lytic induction. We transfected the EBV 2089 producer cells with BZLF1 and BALF4 expression plasmids (to induce optimal virus synthesis) (Hammerschmidt and Sugden, 1988; Neuhierl et al., 2002) together with a plasmid (p7180) that expresses a carboxy-terminal fusion of the gp350 glycoprotein with a codon-optimized bacterial ß-lactamase gene. The chimeric protein, termed gp350:BlaM, was readily incorporated into infectious virions (data not shown) similar to HIV particles that contain a chimeric β-lactamase-Vpr protein (BlaM-Vpr). Upon infection the chimeric protein is delivered into the cytoplasm of HIV target cells as a result of virion fusion (Cavrois et al., 2002; Cavrois et al., 2014; Jones and Padilla-Parra, 2016).

We generated wt/B95.8 (2089) virus stocks with gp350:BlaM protein and subsequently infected PBMCs as illustrated in Figure 7A for four hours. Then, cells were loaded with a cell permeable substrate CCF4-AM and analyzed by flow cytometry. The substrate CCF4-AM is composed of a hydroxycoumarin donor conjugated to a fluorescein acceptor via a β-lactam ring (Jones and Padilla-Parra, 2016). When target cells are infected with gp350:BlaM assembled EBV and β-lactamase reaches the cytoplasm, the enzyme cleaves the β-lactam ring of the CCF4-AM substrate. Cleavage causes a shift of the emission wavelength from 520nm (green) to 447nm (blue), which can be easily detected by flow cytometry (Fig. 7A upper panel).

**Figure 7.**
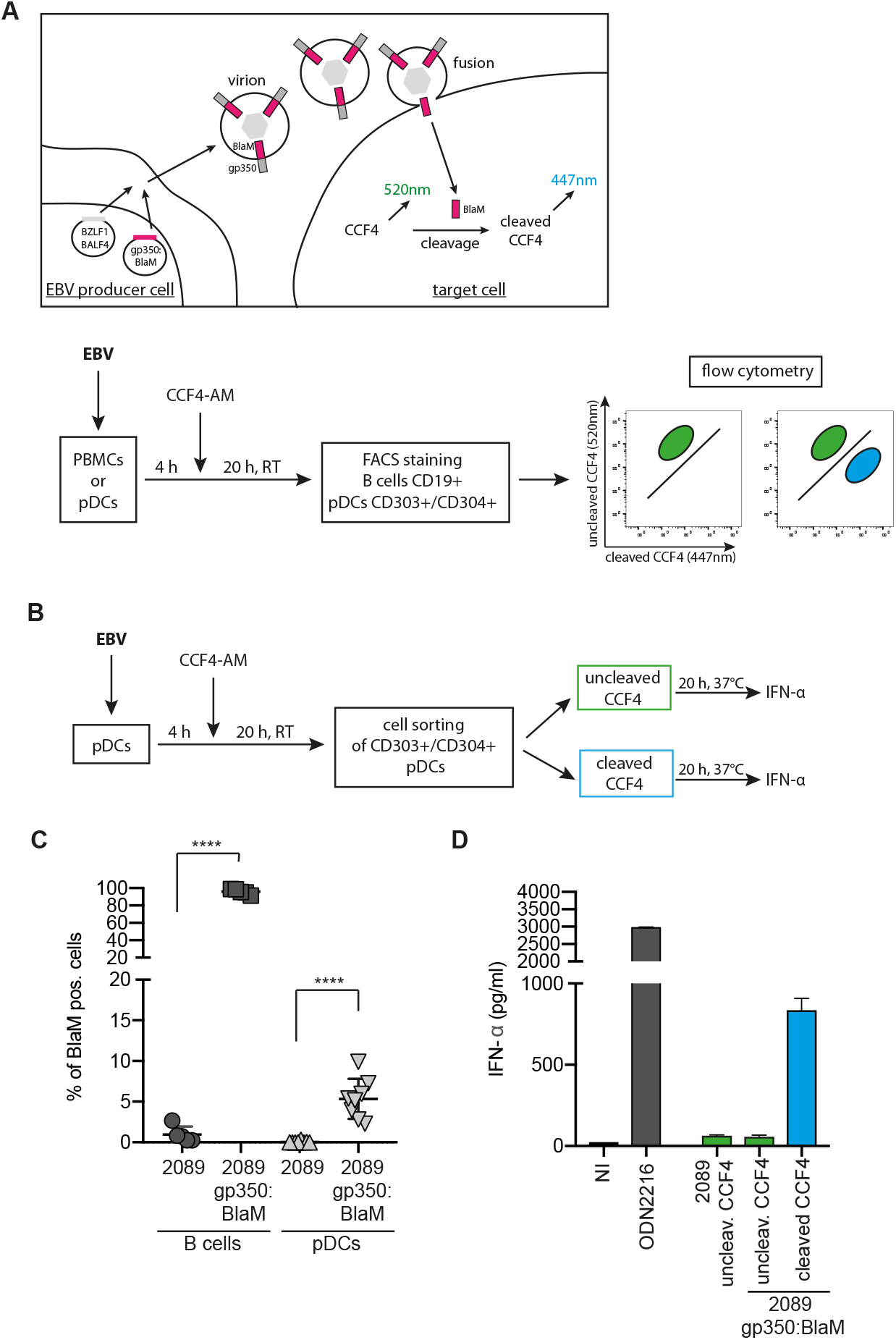
EBV infects a fraction of pDCs which are the main producer of IFN-α. **(A)** Schematic overview of the EBV virion based β-lactamase (BlaM) fusion assay. Expression plasmids encoding BZLF1, BALF4 and the gp350:β-lactamase (BlaM) fusion protein were co-transfected into EBV producer cells (HEK293 wt/B95.8 (2089), Supplementary Table 1). Target cells were infected with the gp350:BlaM assembled virions for four hours and were subsequently loaded with the CCF4-AM substrate. If β-lactamase reaches the cytoplasm of the target cells it cleaves the β-lactam ring of CCF4, which leads to a detectable shift in emission wavelength from 520 nm to 447 nm. Below, a schematic workflow of the steps of the experiment performed in panel C is shown. PBMCs were prepared from human buffy coats and pDCs were isolated via negative selection. PBMCs and pDCs were infected with gp350:BlaM assembled wt/B95.8 (2089) EBV at a MOI of 1. As negative controls the cells were infected with equivalent doses of unmodified wt/B95.8 (2089) EBV. All cells were cultivated for four hours before they were loaded with CCF4-AM and incubated at RT for 20 hours. In case of PBMCs, the cells were stained for B cells using a CD19 antibody and in case of isolated pDCs the cells were stained with two CD303 and CD304 specific antibodies. The percentage of B cells and pDCs with green (uncleaved CCF4 substrate) and blue (cleaved CCF4 substrate) fluorescence was determined using flow cytometry. **(B)** Schematic representation of the steps of the experiment performed in D. pDCs were isolated via negative selection out of PBMCs gained from buffy coats. pDCs were treated with EBV wt/B95.8 (2089) or EBV wt/B95.8 (2089) gp350:BlaM for 4 h. Cells were washed and loaded with CCF4-AM at room temperature for 20 h. The pDCs were sorted for green and blue cell populations and same cell numbers were plated on 96-well cluster plates. The cells were incubated at 37 °C for 20 h and the supernatants were collected. The IFN-α levels were quantified by ELISA. **(C)** Results of experiments with B cells and pDCs as detailed in panel A. The y-axis shows the fraction of cells with blue fluorescence after cleavage of CCF4. Controls are cells infected with unmodified wt/B95.8 (2089) EBV. **(D)** Results of experiments with isolated pDCs obtained from buffy coats as described in panel B. 2×10^4^ pDCs which had been sorted according to their blue fluorescence release close to 1 ng/ml IFN-α, whereas the same number of pDCs that did not show a fluorescence shift in the emission wavelength released very little IFN-α. As negative and positive controls, pDCs that did not show a fluorescence shift were left uninfected or were stimulated with the TLR9 agonist ODN2216.

To analyze if EBV can fuse with pDCs (i.e. infect them), we isolated pDCs from PBMCs and incubated them for four hours with virus stocks of gp350:BlaM assembled EBV wt/B95.8 (2089) or with EBV wt/B95.8 (2089) lacking gp350:BlaM. As an additional negative control we left the cells uninfected. As a positive control, we infected primary B lymphocytes, because they are the main target cells of EBV. After incubating the cells for four hours with the two virus stocks and control samples, the cells were washed, loaded with the β-lactamase substrate CCF4-AM overnight at room temperature and analyzed by flow cytometry (Fig. 7A lower panel). Only when gp350:BlaM assembled EBV successfully infects cells, β-lactamase cleaves the CCF4-AM substrate in the cytoplasm resulting in a shift from green to blue fluorescence.

The main target cells of EBV are B lymphocytes. As expected, almost all B cells in PBMCs turned blue after infection with gp350:BlaM assembled wt/B95.8 (2089) (Fig. 7C). PBMCs infected with EBV lacking gp350:BlaM, uninfected cells or cells treated with gp350:BlaM assembled EVs (which act as a negative control) did not contain any cells with blue fluorescence (Supplementary Fig. 6, upper panel). To investigate whether EBV can also infect pDCs, we isolated pDCs from PBMCs using a negative selection protocol and repeated the BlaM fusion assay. pDCs infected with gp350:BlaM assembled EBV showed approx. 5 % cells with blue fluorescence indicating that a small fraction of pDC fuses with EBV virions (Fig. 7C; Supplementary Fig. 6, lower panel). pDCs incubated with EVs assembled with gp350:BlaM or incubated with EBV lacking gp350:BlaM did not reveal a shift from green to blue cell fluorescence indicating that the novel assay reflects true fusion mediated by infection with EBV (Supplementary Fig. 6). Our newly established EBV BlaM fusion assay documents that EBV can infect pDCs although at a rather low level compared with the virus’s cognate B cells.

Finally, we asked whether the pDCs belonging to the small fraction that EBV infects are the main responders and thereby, main releasors of type I IFN upon viral uptake and fusion. We infected pDCs enriched from PBMCs with gp350:BlaM assembled wt/B95.8 (2089) EBV for four hours, loaded the cells with CCF4-AM and sorted them for green (non-infected) or blue (infected with gp350:BlaM assembled EBV) populations. Identical numbers of sorted green or blue cells were plated in 96-well cluster plates which were incubated at 37°C overnight. The next day, the supernatants were analyzed for IFN-α concentrations by ELISA. Sorted blue cells, which had been infected with gp350:BlaM assembled EBV showed 15 fold higher IFN-α levels than sorted non-infected cells with green fluorescence (Fig. 7D). When we infected the same number of unsorted pDCs with an identical dose of wt/B95.8 (2089) EBV their supernatant contained low IFN-α levels comparable to the sorted, green pDCs incubated with gp350:BlaM assembled wt/B95.8 (2089) EBV (Fig. 7D). This result was expected given the low prevalence of infected cells in unsorted pDC populations. The data indicate that those pDCs that EBV can infect respond with massive IFN-α release, whereas cells that presumably also take up EBV but degrade the incoming virions in the lysosomal pathway contribute little to IFN-α synthesis.

In summary, our experiments reveal that EBV inefficiently fuses with pDCs and that these infected pDCs are the main producers of INF-α. The IFN-α secretion process depends on the presence of unmethylated EBV DNA, which is sensed by the PPR TLR9. Furthermore, the EBV-encoded miRNAs regulate the expression of genes involved in type I IFN secretion and genes involved in the response to type I IFNs in B cells in the pre-latent phase. In pDCs cells, viral miRNAs seem to modulate cellular genes of the type I IFN pathway, but depending on the composition of the different virus preparations only to a minor extent. Unexpectedly, our experiments indicate that the abundantly expressed non-coding small EBERs do not seem to have a role in inducing type I IFN in either B cells or pDCs.

## Discussion

In this study, we examined early anti-viral responses in primary B-lymphocytes and pDCs infected with EBV and EBV mutants. We found that pDCs are the main source of immediate IFN-α release whereas newly infected B cells release comparably low levels of type I IFNs later in the early, pre-latent phase of infection. In infection experiments with wild-type and mutant EBVs, we determined that viral miRNAs interfere with the secretion of pro-inflammatory cytokines and IFN-α from newly infected B cells as well as pDCs. Bioinformatic approaches and data mining of our experiments on established EBV-infected B cell lines revealed many cellular candidate genes that are regulated by EBV-encoded miRNAs and are involved in type I IFN secretion and response pathways. Their validation in subsequent analyses confirmed certain already known mRNA targets but added many new cellular genes in the type I upstream and downstream IFN pathways that appear to be governed by EBV’s numerous viral miRNAs (Fig. 8).

**Figure 8.**
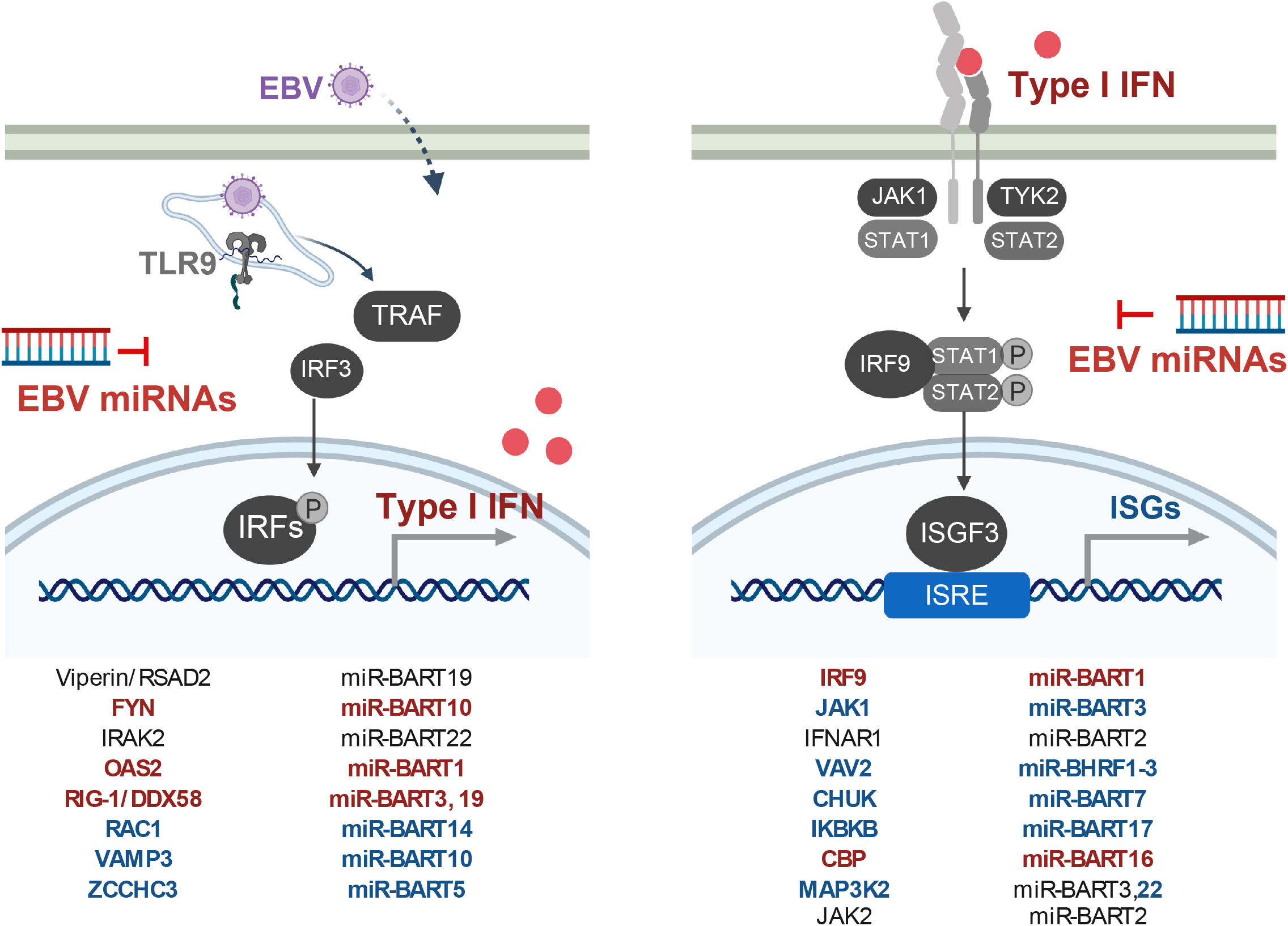
Overview of the identified genes involved in innate sensing and type I IFN response regulated by EBV-encoded miRNAs. **(Left)** The identified target genes regulated by corresponding EBV-encoded miRNAs involved in the regulation of type I IFN production are shown. **(Right)** The identified target genes regulated by corresponding EBV-encoded miRNAs involved in the response to type I IFN are shown. Highlighted in red are targets for which protein expression was affected by loss of the EBV miRNAs; highlighted in blue are targets for which interactions were captured by Ago PAR-CLIP.

Whereas EBV readily infects primary B-lymphocytes, its prime target cells, it was uncertain whether the virus can also infect pDCs, which respond by releasing type I IFN. To address this question and to clarify this so far controversial topic, we developed a novel assay based on viral delivery of an enzymatic activity, ß-lactamase, which we fused to gp350, an abundant viral component in the membrane of EBV virions. We found that EBV can readily fuse with a small fraction of pDCs, which, as a consequence, release large amounts of IFN-α compared to the fraction of pDCs that do not fuse with EBV. We also document that pDCs type I IFN release was unequivocally dependent on viral DNA, which is sensed by the endosomal TLR9 receptor.

Our analysis made use of several mutant EBVs to address the contribution of the abundantly expressed small non-coding EBERs and the role of the viral LF2 gene in EBV target cells (Wu et al., 2009). Very surprisingly and contrary to previous publications, deletion of EBV EBERs had no measurable impact on IFN-α release from PBMCs, pDCs, or B-lymphocytes infected with EBV. The cytoplasmic and endosomal pattern recognition receptors RIG-I and TLR3, respectively, have been reported to become activated by EBERs, inducing type I IFN release (Samanta et al., 2006); Iwakiri et al., 2009, J Exp Med, 206, 2091-9}. EBERs were also reported to confer resistance against IFN-α induced apoptosis by binding to PKR and inhibiting its phosphorylation (Nanbo et al., 2002). In light of these publications, one would expect a reduced type I IFN release in infection experiments with EBER deleted mutant EBVs, which was not the case (Fig. 5). Our experiments differed from these published studies that used established B cell lines, mostly Burkitt lymphoma cells, and experimental settings that required the ectopic expression of EBER molecules. In contrast, we infected and analyzed primary human cells and concluded our observations within a couple of days after infection. Similarly to EBV mutants devoid of EBERs, deletion of the viral gene coding for LF2 resulted in no measurable change of IFN levels compared with infections with r_wt/B95.8 EBV (Fig. 5). This result is conflicting with the literature describing LF2 as a type I IFN antagonist (Wu et al., 2009). Again, experimental conditions differ substantially as the authors relied on ectopic expression of LF2 in luciferase assays and protein-protein interactions with IRF7 in 293T cells (Wu et al., 2009).

Our findings with B cell infection experiments (Fig. 2) provided a conundrum as these results excluded the non-coding EBERs as viral factors that cause the release of pro-inflammatory cytokines (Fig. 2A,B) and IFN-α (Fig. 2C) and which the many viral miRNAs regulate. As a consequence, unknown viral PAMPs likely exist and trigger RIG-I or alternative pattern recognition receptors that presumably recognize them. Recently, several KSHV RNA fragments were identified to be sensed by RIG-I in a RNA Pol III independent manner (Zhang et al., 2018). It seems plausible that also in the context of EBV infection certain viral RNAs other than EBERs might trigger RIG-I or other pattern recognition receptors in infected B cells. The situation in this cell type differs from experiments with PBMCs and pDCs where we could identify virion DNA as the culprit and inducer of IFN-α release (Fig. 6) confirming previous reports (Lim et al., 2007; Fiola et al., 2010; Quan et al., 2010; Severa et al., 2013). Virion DNA is virtually free of methylated cytosine residues very similar to prokaryotic DNA due to a unique strategy of viral DNA replication during the productive, lytic phase of EBV infection (Buschle and Hammerschmidt, 2020).

We established a novel BlaM assay to study the potential of EBV to infect cell populations other than primary B-lymphocytes. The BlaM fusion assay (Fig. 7) identified a minor fraction of pDCs that EBV infects and which releases much higher IFN-α levels than pDCs that were not detectably infected (Fig. 7D). The assay monitors the first steps in viral infection – adhesion, endosomal uptake and fusion of cellular and viral membranes. It makes use of a transmembrane protein which, upon fusion, translocates to the cytoplasmic compartment where the prokaryotic ß-lactamase moiety cleaves its ingenious substrate (Cavrois et al., 2002; Cavrois et al., 2014). Coupling the ß-lactamase reporter domain to gp350 (or another type I transmembrane protein) prevents the secretion of free enzyme and its spontaneous uptake by other cells (Albanese et al., 2020). Consequently, the BlaM assays is free of erroneous background signals supporting the identification of only positive cells in a large population of non-infected cells (Fig. 7C,D) or to detect low level or inefficient fusion of extracellular vesicles with recipient cells (Albanese et al., 2020). The BlaM assay records fusion, only, and is not informative as to whether viral infections are abortive, productive or lead to latent infection. As the assay does not rely on *de novo* gene expression, it can identify cells that are infected but are resistant to subsequent viral gene expression and replication. These cells might still present epitopes of the incoming viral particle to immune cells such as CD4+ T cells contributing to beneficial or adverse immune responses of the infected host organism.

Previously published data show conflicting results regarding pDCs and EBV infection. Fiola et al. postulated that EBV does not establish infection in pDCs but responded to EBV particles or isolated EBV DNA similar to our findings (Fiola et al., 2010). On the other hand, certain isolated EBV strains efficiently infected monocytes which had been stimulated with GM-CSF and IL-4 to induce their differentiation to DCs (Guerreiro-Cacais et al., 2004). In that study, the efficiency of EBV entry into monocytes was monitored by measuring green fluorescence intensity after 48 hours. In our fusion assay, we evaluated the immediate fusion of EBV with pDCs as our incubation period was just four hours before loading the cells with the CCF4-AM substrate, but we found only a small fraction (< 10 %) to undergo fusion (Fig. 7C). In contrast, efficient infection of pDCs through viral binding to the MHC class II molecule HLA-DR and expression of EBV-encoded genes was reported (Severa et al., 2013) together with MHC class II mediated uptake, degradation and subsequent activation of TLR9 followed by IFN-α secretion (Quan et al., 2010). In contrast, we did not observe expression of viral genes (or GFP encoded in the backbone of the viral genomic DNA) in pDCs even when the cells were incubated for up to two days after infection with EBV (data not shown). EBV infection of pDCs and monocytes was reported to reduce biological activities of these cell types (Li et al., 2002; Gujer et al., 2019). Li et al. further postulated that pDCs are not infected, but apoptotic and infected B cells cause the presentation of EBV antigens on MHC class I molecules of DCs. Recently it was reported that pDCs are able to undergo trogocytosis, i.e. to conjugate with antigen presenting cells and to extract surface molecules, including MHC class I molecules, presenting them on their own cell surface. With this mechanism non-infected pDCs could be able to present EBV specific epitopes on MHC class I molecules (Bonaccorsi et al., 2014).

Prior to our study, an accurate and direct method to monitor EBV infection of different cell populations was not available. Our newly introduced gp350-BlaM fusion assay is based on the uptake of EB virions and the fusion of their viral membrane with cellular, possibly endosomal membranes such that the content of the virion is released into the cytoplasm of EBV’s host cell. As such, the method can monitor the very first steps of virus entry and fusion and does not depend on subsequent events such as viral *de novo* transcription for example. Thus, our novel approach can precisely determine the fraction of cells, with which EBV virions fuse within hours. Using this assay, we demonstrate that EBV virions containing gp350-BlaM fuse with only a small fraction of pDCs, approximately 5 % but with the vast majority of primary B-lymphocytes (Fig. 7C). Interestingly, among the abundance of DCs in general, only the small number of pDCs acts upon EBV infection by releasing IFN (Gujer et al., 2019). It also seems as if a so far unknown subpopulation of pDCs might exist that EBV targets (Fig. 7C,D), but their cellular identity is uncertain.

A closer look at the literature shows that there is a network of identified EBV-encoded miRNAs and their regulated targets. The identified target genes are mainly located in adaptive and innate immunity and immune escape mechanisms (Albanese et al., 2017), what also corroborates our hypothesis that miRNAs interfere with type I IFN signaling as part of the innate immune escape during EBV infection (Gallo et al., 2020; Iizasa et al., 2020; Li et al., 2020). With our experiments shown in Figures 3 and 4 and documented in Supplementary Figures 3 and 4, we confirmed a number of host cell transcripts targeted by viral miRNAs, but we also identified several previously unknown effectors and their cellular transcripts (Fig. 8). Already in 2017, through gene expression profiling experiments, miR-BART6-3p was reported to regulate RIG-I and IFN-β responses (Lu et al., 2017). Also, in our study we could confirm that RIG-I is a target of EBV-encoded miRNAs in dual luciferase assays and in recently infected B cells. We observed that miR-BART3 and miR-BART19 are both able to regulate the RIG-I 3’UTR (Fig. 3A). These findings indicate that possibly a combination of different miRNAs during EBV infection leads to a regulation of the same cellular gene. *CBP*, which we evaluated to be regulated by miR-BART16 (Fig. 4B), was already reported to be target of this viral miRNA (Hooykaas et al., 2017).

*CBP, RIG-I, FYN, IRF9*, and *JAK1* are good examples of single genes central to IFN signaling and regulated concomitantly by several EBV-encoded miRNAs (our study, Hooykaas et al., 2017; Cox et al., 2015). On the contrary, there are also certain genes that were reported to be regulated by EBV miRNAs, which we could not confirm in 3’UTR luciferase reporter assays. In fact, the majority of candidates from our initial bioinformatics analysis did not pass our criteria as targets when analyzed in dual luciferase reporter assays. For each candidate, we cloned the 3’UTR as well as introduced point mutations into the suspected seed match sequence to ascertain whether a functional biochemical interaction between the miRNA and its predicted mRNA target could occur. For most candidates of the screen, we were unable to confirm that the predicted miRNA interaction sites were functional. However, we found that certain cellular proteins are clearly upregulated in r_ ΔmiR-infected B cells, such as TLR7 and TLR9 (Fig. 3F) but, in luciferase reporter assays, we failed to identify any viral miRNAs that could regulate the 3’UTRs of these two transcripts (Supplementary Fig. 7). Thus, TLR7 and TLR9 do not appear to be direct targets of EBV miRNAs, but their expression levels are certainly impacted by the presence of the viral miRNAs in EBV infected cells.

Key experiments in this study rely on phenotypes in primary immune cells with an EBV strain and its mutant derivatives that are based on the reconstituted wildtype EBV strain r_wt/B95.8 as shown in Figure 1. This EBV strain encodes all 25 pre-miRNAs that are processed to 44 mature viral miRNAs presumably expressed at physiological levels in EBV infected cells. Recently, Pich et al. showed that the deletion of all miRNAs in r_ΔmiR leads to an increase in apoptosis of EBV infected primary B cells, as well as a lower division index and reduced numbers of cells presented in S-phase demonstrating that EBV-encoded miRNAs are involved in the regulation of cell cycle functions (Seto et al., 2010; Pich et al., 2019). Furthermore, studies in primary B cells showed that viral miRNAs can interfere with their immune recognition by antiviral CD4+ and CD8+ T cells (Albanese et al., 2016; Tagawa et al., 2016). Using EBV mutant derivatives (Fig. 1), we identified an additional role of viral miRNAs in governing type I IFN responses upon infection, but we were puzzled to find that the abundant EBERs do not have an apparent inducing function in this pathway. On the contrary, their deletion caused a slight but consistent increase in IFN in cells infected with the double mutant ΔEBER/ΔmiR suggesting that the EBERs might even have a damping effect on the release of type I IFN in PBMCs and pDCs (Fig. 5A,C). Clearly, this finding warrants further exploration both in infection experiments and at the molecular level of type I IFN release.

## Materials and Methods

### Construction of mutant EBVs

All recombinant EBVs used in this study are based on the maxi-EBV plasmid p2089, which contains the B95-8 EBV genome (Delecluse et al., 1998). The EBV strain B95-8 has a deletion of several pre-miRNA loci, lacks the second lytic origin of DNA replication (*oriLyt*) and several other genes, which are present in EBV field strains. As described in Pich et al. a new wildtype-like strain based on B95-8 expressing all EBV-encoded miRNAs at physiological level was established. This newly generated reconstituted wildtype strain is called r_wt/B95.8 (6008) and was constructed by introducing a DNA fragment of the EBV strain M-ABA repairing the deletion in B98-5 and the maxi-EBV plasmid p2089 (Pich et al., 2019). Furthermore, parts of the F-plasmid backbone were replaced by introducing an artificial open reading frame encoding eGFP and *pac (puromycin N-acetyltransferase* conferring puromycin resistance). This new r_wt/B95.8 (6008) strain expresses all viral miRNAs at physiological levels, carries the second *oriLyt* and expresses LF1, LF2 and LF3 viral proteins, which are absent in the B95-8 EBV genome. A second EBV strain used in this study is the r_ΔmiR (6338) strain lacking all of the EBV-encoded pre-miRNA loci. It was generated as described in Pich at al. (Pich et al., 2019). Furthermore, the genes of the two viral non-coding EBER RNAs were deleted in r_wt/B95.8 and r_ΔmiR to obtain mutant EBVs termed ΔEBER and ΔEBER/ΔmiR, respectively (Fig. 1) (Pich et al., 2019). Based on genome of r_wt/B95.8 (6008) the viral LF2 gene was disabled by introducing a stop codon in the coding sequence of the LF2 gene. This EBV mutant was called ΔLF2 (Fig. 1, Supplementary Tab. 1).

### Cells and culture

Cells used in this study were mostly cultivated in RPMI 1640 medium supplemented with 8 % fetal calf serum (FCS), 100 μg/ml streptomycin, 100 U/ml penicillin, 1 mM sodium pyruvate, 100 nM sodium selenite and 0.43 % α-thioglycerol at 37 °C and 5 % CO_2_. HEK293 2089 cells were cultured with the addition of 100 μg/ml hygromycin and HEK293 6008, 6338, 6431, 6432 and 6522 cells with the addition of 500 ng/ml puromycin. 293T cells for dual luciferase reporter assays were cultured in DMEM medium supplemented with 8 % FCS, 100 μg/ml streptomycin, 100 U/ml penicillin and 1 mM sodium pyruvate (Fig. 3A-E, Fig. 4B-E, Supplementary Fig. 3A-C). HEK293T cells were maintained in high glucose DMEM supplemented with 10% fetal bovine serum (FBS) and 1% penicillin/streptomycin/glutamine (Supplementary Fig. 3D-K). DLBCL cells (IBL1, IBL4, and BCKN1) were maintained in RPMI supplemented with 10% FBS and 1% penicillin/streptomycin/glutamine.

### Preparation and quantification of infectious viral stocks

HEK293 virus producer cell lines were established after transfection of the different maxi-EBV plasmids and subsequent selection with puromycin. To generate virus stocks clonal producer cells were transfected using PEI Max with the two plasmids, p509 and p2670, to induce the lytic cycle of EBV. p509 encodes the BZLF1 gene, which triggers lytic phase reactivation (Hammerschmidt and Sugden, 1988) and p2670 the BALF4 gene, which increases the infectivity of the recombinant EBVs (Neuhierl et al., 2002). Three days post transfection supernatants were collected and centrifuged for 10 min at 1,200 rpm and 30 min at 5,000 rpm to remove cell debris. To determine the titers of the different virus stocks we used Raji cells as described (Steinbrück et al., 2015). In detail 1×10^5^ Raji cells were incubated with different amounts of virus stocks in a volume of 1 ml for three days at 37 °C. By flow cytometry the percentage of GFP positive cells was determined and the linear regression equation was calculated as described (Steinbrück et al., 2015). To concentrate the virus, a further ultracentrifugation step with an iodixanol (Optiprep) cushion was used. Therefore, the virus supernatants were loaded on 2-3 ml iodixanol (5 volumes of solution A (iodixanol) and 1 volume of solution B (0.85% (w/v) NaCl, 60 mM Hepes, pH 7.4) and centrifuged for 2 h at 28,000 rpm at 4 °C. Concentrated virus in the interphase was transferred to microfiltration tubes (Amicon Ultra-15) and centrifuged at 2000 rpm at 4 °C. Concentrated virus stocks were quantified using Raji cells as described above.

### Cloning and construction of luciferase reporter plasmids

Human 3’UTR sequences were amplified from genomic DNA by PCR with primers harboring Gateway-compatible extensions or restriction enzyme sites. RIG-I (DDX58), Viperin (RSAD2), IKKβ (IKBKB), IRAK2, FYN, CBP, IRF9, JAK1, OAS2, JAK2 and MAP3K2 are cloned downstream of Renilla luciferase (Rluc) in the modified psiCHECK-2 (Promega) plasmid p5522 using the Gateway cloning technology (m Fisher Scientific) (Fig. 3A-E, Fig. 4B-E, Supplementary Fig. 3A-C). To generate other 3’UTR luciferase reporters, sequences were PCR amplified from genomic DNA of EBV-infected B cells and cloned into the XhoI and NotI sites downstream of *Renilla* luciferase in the psiCheck2 dual luciferase reporter vector containing an expanded multiple cloning site. When achievable, the entire 3’UTR was cloned for a given target. For longer 3’UTRs, a minimum of 1 kb containing the region predicted to be targeted by each miRNA was cloned (Supplementary Fig. 3D-K). EBV miRNA expression vectors in pLCE are previously described (Skalsky et al., 2014).

### Luciferase reporter assays

HEK293T cells were co-transfected with 20 ng of sis-inducible element (SIE) firefly luciferase reporter, 250 ng of control vector (pLCE) or EBV miRNA expression vector, and 20 ng of pRL-SV40 renilla luciferase reporter as internal control, using Lipofectamine 2000 (Thermo Fisher) according to the manufacturer’s instructions. 48 hrs post-transfection, cells were mock treated or treated with 10 unit/mL of uIFN for 6 hrs and collected in 1× passive lysis buffer (Promega). Lysates were assayed for luciferase activity using the Dual Luciferase Reporter Assay System (Promega) and a luminometer with dual injectors. All values are reported as relative light units (RLU) relative to luciferase internal control and normalized to mock treated pLCE control vector (Fig. 4F).

The 3’UTR reporter plasmids were co-transfected into HEK293T cells using PEI max together with the indicated pCDH-EF1-MCS expressing plasmids (System Biosciences) containing a cloned pre-miRNA gene of interest. 24 h post-transfection the luciferase activity was measured using the Dual-Luciferase Assay Kit (Promega) and an Orion II Microplate Luminometer (Titertek-Berthold). The activity of renilla luciferase (Rluc) was normalized to the activity of firefly luciferase (Fluc) expressed by the psiCHECK-2 reporter. Site-directed mutagenesis of putative EBV miRNA seed match sites in the reporter plasmids (Supplementary Fig. 2) was performed as described previously (Tagawa et al., 2016) (Fig. 3A-E, Fig. 4B-D, Supplementary Fig. 3A-C).

HEK293T cells plated in 96-well black-well plates were co-transfected with 20 ng of 3’UTR reporter and 250 ng of control vector (pLCE) or miRNA expression vector (Skalsky et al., 2014) using Lipofectamine 2000 (Thermo Fisher) according to the manufacturer’s instructions. 48-72 hrs post-transfection, cells were collected in 1× passive lysis buffer (Promega). Lysates were assayed for luciferase activity using the Dual Luciferase Reporter Assay System (Promega) and a luminometer with dual injectors. All values are reported as relative light units (RLU) relative to luciferase internal control and normalized to pLCE control vector (Supplementary Fig. 3D-K).

### Generation of gp350:BlaM assembled extracellular vesicles (EVs) and their quantitation

To assemble extracellular vesicles (EVs) with gp350:BlaM we transfected HEK293 cells with the p7180 plasmid, which codes for a fusion of gp350 and a codon-optimized BlaM (gp350:BlaM). Three days after transfection supernatants were harvested as described above and EVs were sedimented by ultracentrifugation at 24,000 rpm for 2 h. The EV stocks were quantified using the Elijah assay. 2×10^5^ Elijah cells (Rowe et al., 1985) were incubated with different volumes of EVs or a calibrated wt/B95.8 (2089) reference EBV stock with a known virus concentration (as measured in GRU per milliliter). The Elijah cells were incubated at 4 °C for 3 h on a roller mixer, subsequently washed twice with PBS and resuspended in 50 μl staining buffer (PBS, 0.5% BSA, 2 mM EDTA) containing 1 μl of the anti-gp350 antibody 6G4 coupled to Alexa Fluor 647. Cells were incubated at 4 °C for 20 min, washed with staining buffer and analyzed by flow cytometry. The mean fluorescence intensity (MFI) values of gp350 were recorded and the concentration of the EVs was estimated using the linear regression equation obtained with the aid of the reference EBV stock.

### Isolation of B cells from adenoids and infection

Adenoid biopsies were washed with PBS and mechanically disintegrated with two sterile scalpels in a sterile petri dish. The cell suspension was filtered through a 100 μm mesh cell strainer. To obtain a maximum of single cells this procedure was repeated. The cell suspension was brought to 50 ml with PBS, mixed and centrifuged at 300 g for 10 min. The cell pellet was resuspended in 30 ml PBS and 0.5 ml defibrinated sheep blood for T cells rosetting was added. The cell suspension was layered onto 15 ml Ficoll Hypaque and centrifuged at 500 g for 30 min. The cells were carefully collected from the interphase and transferred to a new 50 ml tube. The cells were washed two times with PBS, resuspended in 800 μl staining buffer (PBS, 0.5% BSA, 2 mM EDTA) and 70 μl CD3-APC beads were added for 15 min at 4 °C. A depletion of CD3 positive cells was done by magnetic bead sorting. The cells were washed in 10 ml staining buffer and resuspended in pre-warmed medium. After counting the cells were infected with titered virus stocks at a density of 1 ×10^6^ cells/ml. The next day the cells were centrifuged, resuspended in fresh medium at the same density and cultivated. Four days later (on day five post-infection) the cells were counted using calibrated APC beads as volume standard using a flow cytometer. The cells were incubated for 4 subsequent days at 37 °C at a density of 7×10^5^ cells/ml in a total volume of 2 ml in 24-well cluster plates. Afterwards, supernatants were collected for ELISA analysis.

### Protein lysates from infected B cells and Western blot immunostaining

To determine the protein expression in infected B cells, B cells were isolated as described above. Infection of the isolated cells with the indicated virus stocks was performed at an initial density of 1×10^6^ cells/ml. After 24 hours the cells were centrifuged and resuspended in fresh medium at the same density. Cells were incubated at 37 °C for 4 or 8 additional days. Cells were washed in cold PBS and resuspended in RIPA lysis buffer complemented with protease and phosphatase inhibitors. Cell lysates were frozen at −80 °C. After thawing on ice, the lysates were mixed and centrifuged at 13,000 rpm for 10 min at 4 °C. Supernatants were collected and the protein amount was determined using the Bradford assay. Protein concentrations of the lysates were adjusted using RIPA lysis buffer, Lämmli buffer was added and identical protein amounts of the different samples were loaded on TGX Stain-free Precast gels purchased from Biorad. After the run the gels were activated by a 45 sec UV exposure and electroblotted onto nitrocellulose membranes. The membrane was blocked and incubated with the indicated primary antibodies (Fig. 3F and 4E). The following antibodies were used directed against IPO7 (Sigma, # SAB1402521, 1:1000), Viperin/RSAD2 (Abcam, #ab107359, 1:100), TLR9 (Santa Cruz, # sc-47723, 1 :200), IFIT1 (abcam, #ab70023, 1:1000), IRF-9 (Cell signaling, #28492, 1:1000), TLR7 (Abcam, #ab124928, 1:1000), FYN (Biolegend, #Fyn-59, 1:1000), CBP (Abcam, #ab2832, 1:1000), RIG-I/DDX58 (Adipogen, #AG-20B-0009, 1:1000), and OAS2 (Santa Cruz, #sc-374238, 1:1000). The following secondary antibodies were used: anti-mouse HRP (Cell signaling, #7076S, 1:10000), anti-rabbit HRP (Cell signaling, #7074S, 1:20000). The membranes were scanned using the ChemiDoc Imaging sytem (Bio-Rad) and the images were analyzed and the signals quantitated after total cell protein normalization using the Image Lab 6.0.1 software (Bio-Rad).

### ELISA

For detection of cytokine secretion by B cells, PBMCs or pDCs we used the Human ELISA development Kits from Mabtech for IFN-α pan (#3425-1A-20), IL-12 (#3455-1A-6), IL-6 (#3460-1A-6) and IL-10 (#3430-1A-6). The assays were performed as described in the manufacturer’s protocol using 60-80 μl of cell supernatants.

### Isolation and infection of PBMCs and pDCs

PBMCs were isolated from fresh whole blood samples or buffy coats. The blood was diluted 1:3 or more in PBS and 30 ml of the diluted blood sample was layered onto a 15 ml Ficoll Hypaque cushion in a 50 ml tube. The tubes were centrifuged at 1,850 rpm at 10 °C for 30 min. PBMCs were carefully collected from the turbid interphase and were transferred to a fresh tube. Cells were washed with PBS three times at decreasing centrifugation parameters (1,600 rpm, 1,400 rpm, 1,200 rpm for 10 min, 10 °C). The cell pellet was resuspended in fresh medium and the cells were counted. For the isolation of pDCs from PBMCs the Plasmacytoid Dendritic Cell Isolation Kit II from Miltenyi Biotec (# 130-097-415) was used. The isolation of pDCs was performed as described in manufacturer’s protocol using MACS sorting with LS columns. For ELISA assays 2×10^5^ PBMCs or 2×10^4^ pDCs per well in 200 μl total volume were seeded in 96-well cluster plates. The cells were infected with the indicated virus strains with different MOIs and the cells were incubated at 37 °C for 20 h.

### Ethics

PBMCs were obtained from volunteer blood donors. Adenoid biopsies were obtained from patients from the Department of Otorhinolaryngology of the Universitätsklinikum Ludwig-Maximilian-Universität München. Biopsies originated from disposed tissues from anonymous donors who underwent routine surgery.

### PAR-CLIP datasets and Ingenuity pathway analysis

EBV B95-8 or wild-type LCLs and EBV+ PEL PAR-CLIP datasets are previously described; Ago PAR-CLIP datasets for AIDS-related DLBCL cell lines (IBL1, IBL4, and BCKN1) were generated as previously described (Gottwein et al., 2011; Skalsky et al., 2012; Majoros et al., 2013; Skalsky et al., 2014). AIDS-related DLBCL cell lines were kindly provided by Dr. Ethel Cesarman (Weill Cornell Medical College) and maintained in RPMI 1640 supplemented with 20 % fetal bovine serum, 2 mM L-glutamine, and antibiotics. Briefly, cells were pulsed for 18 hr with 100 μM 4-thiouridine, cross-linked using UV 365 nm, and Ago-associated RNAs were immunopurified using pan-Ago or Ago2 antibodies. Sequencing of barcoded CLIP samples was performed at the OHSU MPSSR on the Illumina HiSeq-2000 platform. Raw sequencing datasets are available through NCBI SRA under Bioproject (in submission).

Reads were processed using the FASTX-toolkit (http://hannonlab.cshl.edu/fastx_toolkit/), aligned to the human genome (hg19) using Bowtie (-v 3, -l 12, -m 10, --best --strata), and Ago interaction sites were defined using the PARalyzer v1.5 (Corcoran et al., 2011) and PIPE-CLIP (Chen et al., 2014) pipelines. For PARalyzer analysis, reads are mapped to the genome and reads that overlap by at least one nucleotide are placed into groups. Groups are then analyzed for T>C conversions and regions with a higher likelihood of T>C conversions are posited as interaction sites so long as the minimum read depth of three reads per site is met. 3’UTR Ago interaction sites were annotated and scanned *ad hoc* for canonical seed matches (>=7mer1A) to mature EBV miRNAs.

To define major cellular pathways regulated by EBV miRNAs, we used Ingenuity Pathway Analysis (IPA) (Qiagen). 3,976 human genes harboring 3’UTR miRNA interaction sites (>=7mer1A) as identified in PAR-CLIP (EBV+ BC1 cells, EBV B95-8 or wild-type LCLs and EBV+ DLBCL cells) were queried. Enriched canonical pathways related to IFN-I signaling were defined by IPA, and individual pathways were selected for visual representation. Genes targeted by EBV miRNAs that modulate these enriched pathways were curated and subject to further validation.

### BlaM fusion assay

To study EBV’s infectivity in different cell types we used the BlaM fusion assay based on studies with recombinant HIV (Cavrois et al., 2002; Cavrois et al., 2014; Jones and Padilla-Parra, 2016). We generated wt/B95.8 (2089) virus stocks using its producer cells, which were co-transfected with p509, p2670 and p7180 plasmid DNAs. p7180 encodes a fusion of gp350 and a codon-optimized β-lactamase. Up to 1×10^6^ PBMCs or isolated pDCs were infected with wt/B95.8 (2089) assembled with gp350:BlaM or with unmodified wt/B95.8 (2089) virus as negative control at 37 °C for 4 h. Cells were washed in 200 μl CO_2_ independent medium (Gibco, #18045-054). The cells were loaded with CCF4-AM substrate (Thermo Fisher, #K1095) in 100 μl staining solution (1 ml CO_2_ independent medium, 2 μl CCF4 (Invitrogen # K1095), 8 μl Solution B (Invitrogen # K1095), 10 μl Probenecid (Sigma, # P8761-25g)) per well and incubated in a humidity chamber in the dark at room temperature overnight. After incubation the cells were washed twice in FACS buffer (PBS, 0.5% BSA, 2 mM EDTA) and stained with a CD19-APC antibody (BioLegend # 302212) to analyze pDCs for 20 min at 4°C to analyze B cells or with two antibodies directed against CD303(BDCA2)-APC (Miltenyi Biotec, #130-097-931) and CD304(BDCA4)-PE (Miltenyi Biotec # 130-112-045). The cells were washed in FACS buffer (PBS, 0.5% BSA, 2 mM EDTA) and analyzed using a Fortessa flow cytometer (Becton-Dickinson). To measure the cytokine secretion of the different pDC populations, the cells were sorted using a BD FACS Aria III instrument according to the fluorescence of the CCF4-AM substrate and identical numbers of sorted cells were seeded in 96-well cluster plates. The cells were incubated at 37 °C for 20 h. The supernatants were collected and IFN-α ELISA assays were performed as described above.

### Reagents

ODN2088 (Miltenyi Biotech, # 130-105-815)

ODN2087 Ctrl (Miltenyi Biotech, # 130-105-819)

Chloroquine (Sigma Aldrich, C6628)

## Acknowledgements

M.B. received an EMBO long-term fellowship by the European Molecular Biology Organization. R.L.S. and work in her laboratory is supported by NIH grants R01-AI143620 and R00-CA175181. This work was financially supported by grants from the Deutsche Forschungsgemeinschaft (grant number SFB1064/TP A13), Deutsche Krebshilfe (grant number 70112875) and German Center for Infection Research (grant number 07.814) to W.H..

## Author contributions

MB and SV performed most of the experiments, DP, CG, YC and DNF provided additional experimental work, TT, MA, RLS and WH designed the experiments and the scientific concept, and MB, SV, RLS, and WH wrote the paper.

---

**Supplementary Figure 1.**
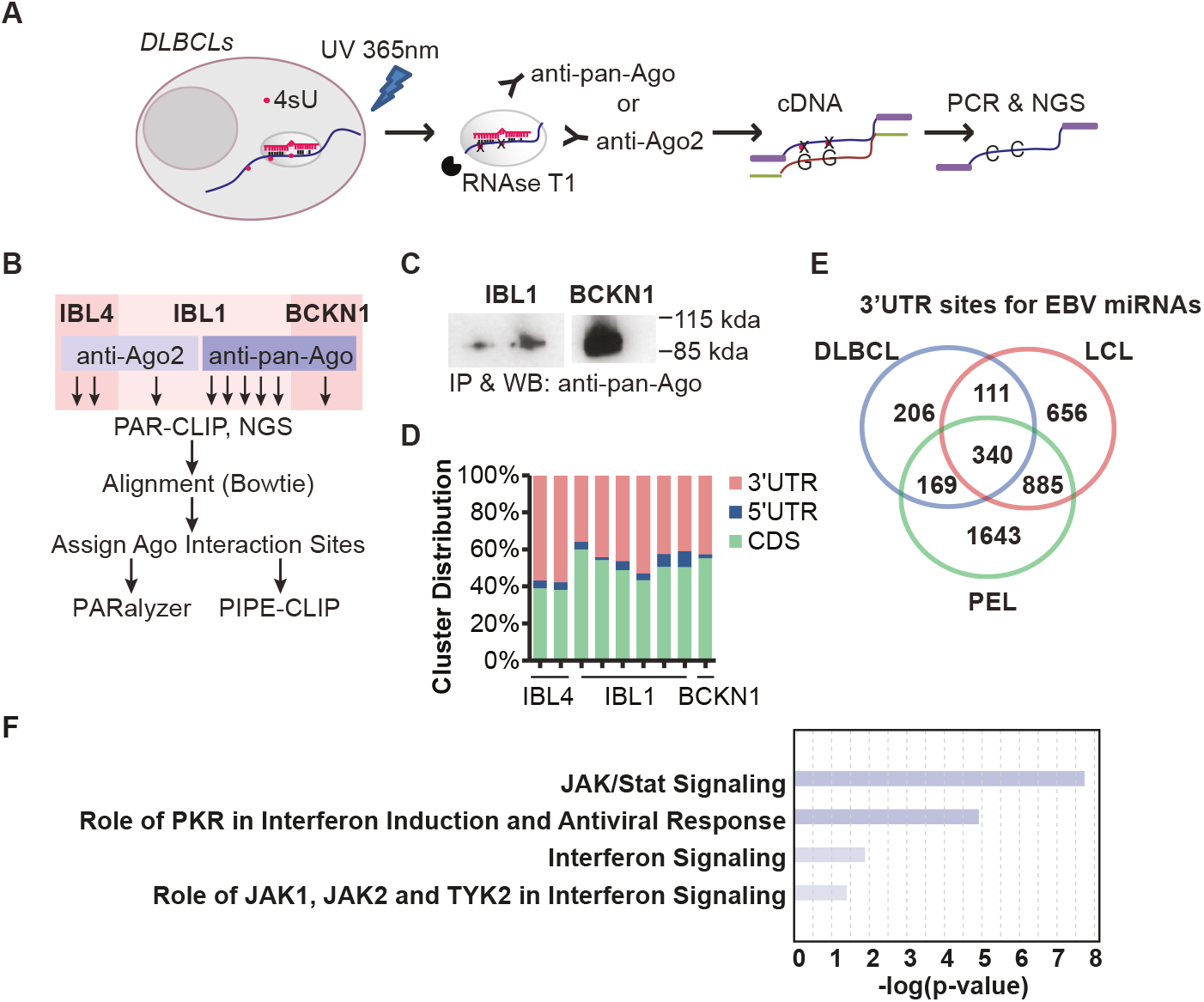
Ago PAR-CLIP analysis identifies viral miRNA targets in EBV-infected B cell lines. **(A)** Overview of PAR-CLIP method. **(B**) Workflow of PAR-CLIP samples analysed and bioinformatics tools utilized to define Ago interaction sites in EBV-infected DLBCL cell lines IBL4 (two replicates), IBL1 (six replicates), and BCKN1 (one replicate). (**C)** Western blot (WB) validation of UV-crosslinked, immunopurified (IP) Ago protein in IBL1 and BCKN1 cells. (**D**) Distribution of Ago interaction sites mapping to human protein-coding transcripts in DLBCL cells. CDS = coding sequence, UTR = untranslated region (**E)** Number of unique 3’UTR sites harbouring canonical seed matches to EBV miRNAs (>=7mer). 340 total 3’UTR sites were commonly identified in Ago PAR-CLIP datasets from DLBCL cells (current study) and published LCL and EBV+ PEL cells. (**F)** Ingenuity pathway analysis of EBV miRNA targets. 3,976 genes identified in PAR-CLIP datasets as EBV miRNA targets (from (E) were queried using IPA core analysis. Enriched canonical pathways related to IFN-I signaling were defined, and individual pathways selected for visual representation.

**Supplementary Figure 2.**
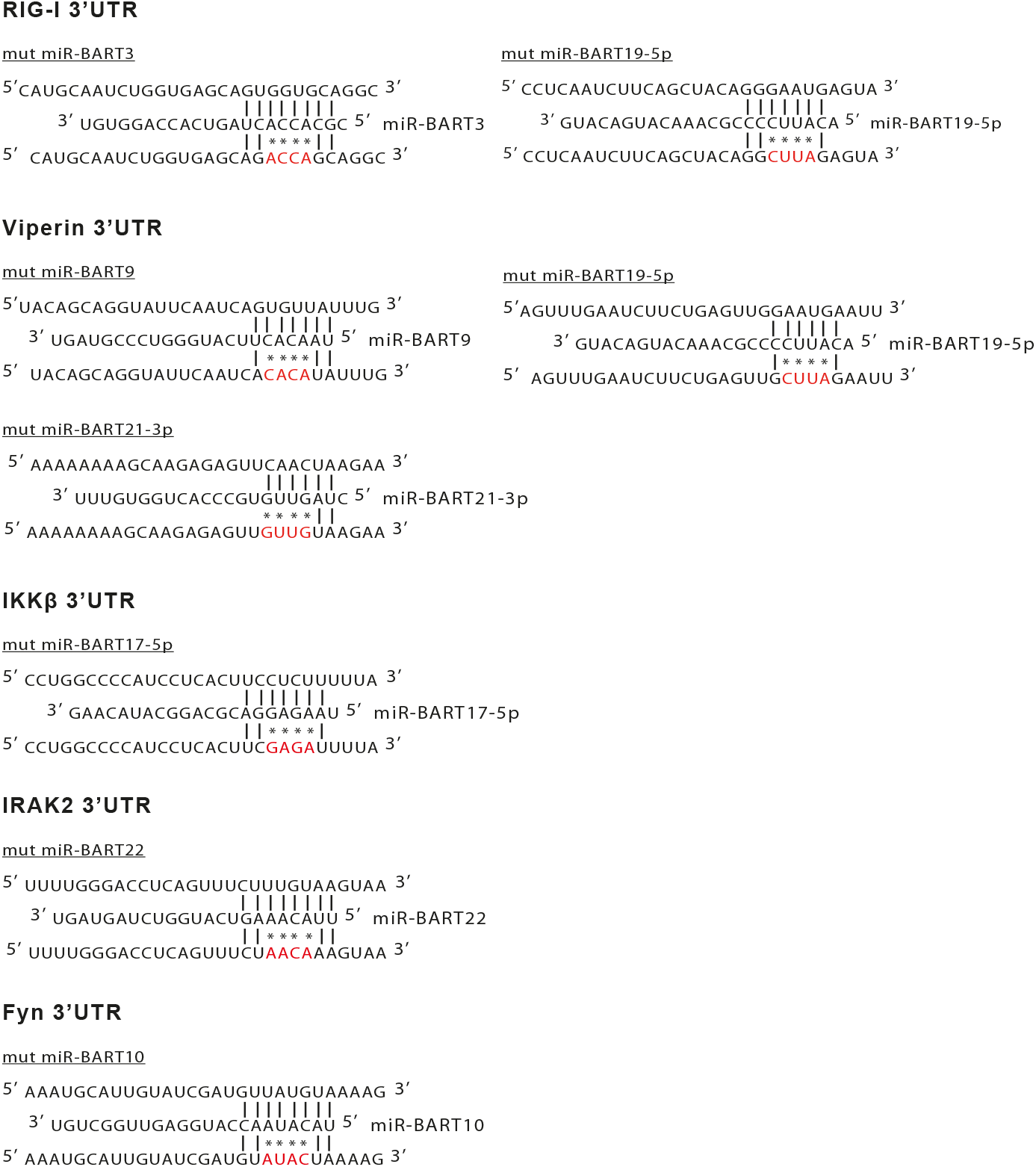
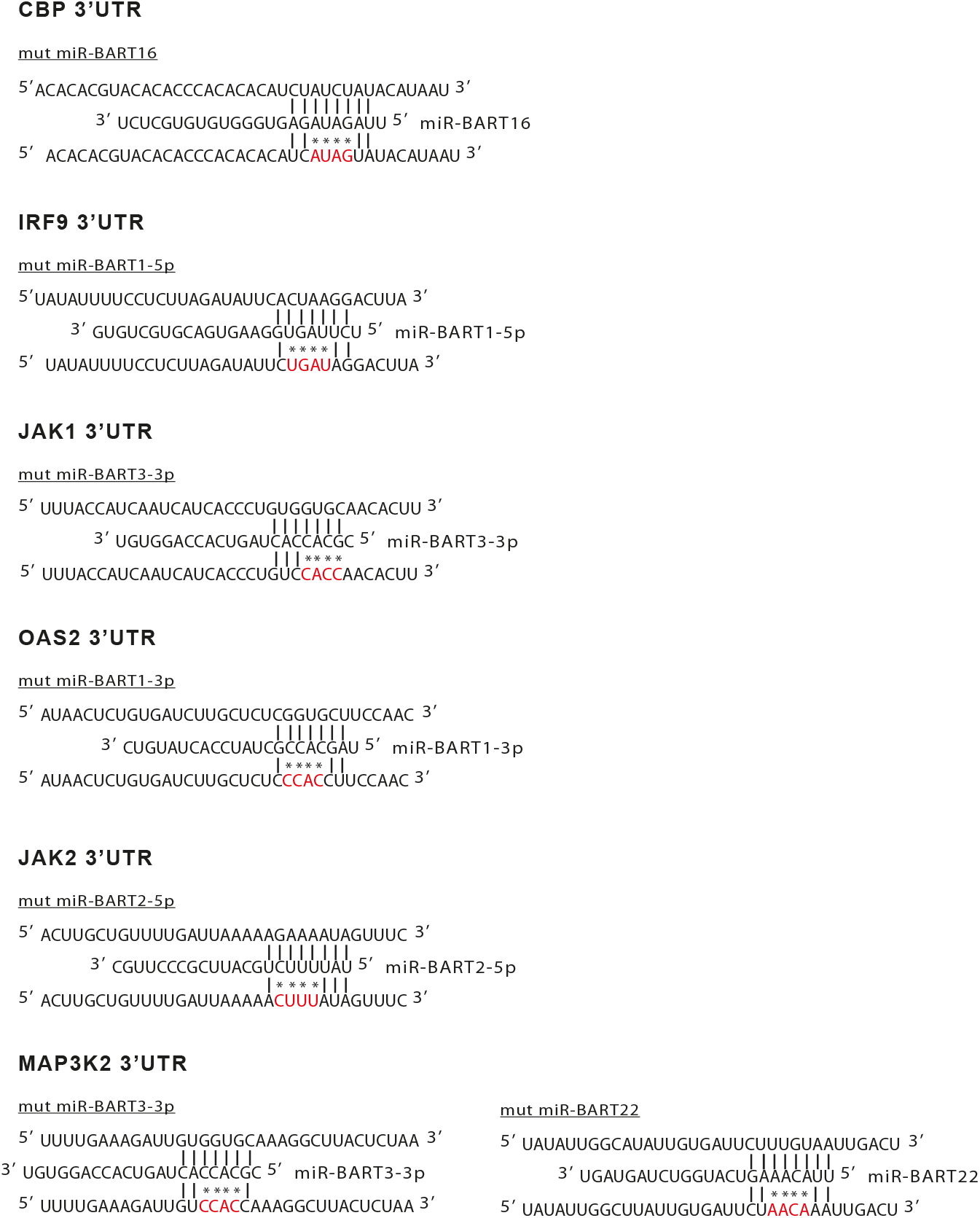
Overview of complementary binding of miRNA seed sequences to 3’UTR target transcripts and point mutations in the mRNAs with presumed seed sequence specificities. The binding sequences of EBV-encoded miRNAs to selected 3’UTR target transcripts of RIG-I/DDX58, IRAK2, IKKβ, Viperin/RSAD2, FYN, OAS2, CBP, IRF9, JAK1, JAK2 and MAP3K2 are shown. The point mutations in the presumed seed sequences are marked in red.

**Supplementary Figure 3.**
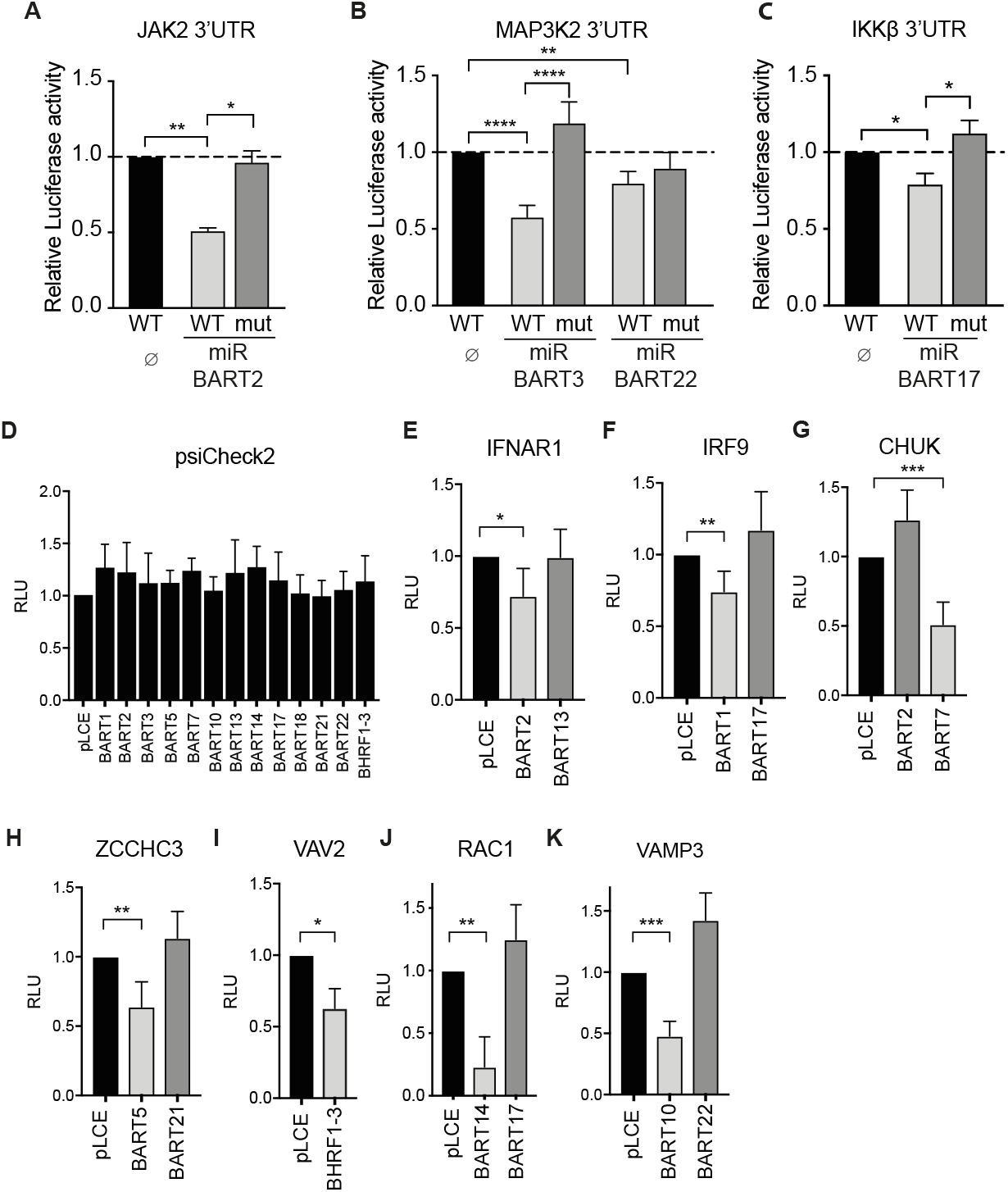
Genes involved in innate sensing and type I IFN response are regulated by EBV-encoded miRNAs. **(A-C)** 293T cells were co-transfected with the indicated wild-type luciferase reporter plasmids (WT) for JAK2, MAP3K2 or IKKβ or reporter plasmids where the predicted seed sequence were mutated (mut) together with or without a miRNA-encoding plasmid. Luciferase expression in these cells was assessed and normalized to lysates from cells co-transfected with the wild-type 3’-UTR reporter and the empty miRNA expression plasmid (∅). P-values were calculated by the one-way ANOVA test. *, P < 0.05; **, P < 0.01; ****, P < 0.0001. **(D-K)** HEK293T cells were co-transfected with psiCheck2 empty vector **(D)** or indicated 3’UTR luciferase reporters **(E-K)** and EBV miRNA expression vectors (pLCE-based). 48–72 hrs post-transfection, cells were lysed and assayed for dual luciferase activity. Reported are the averages of at least four independent experiments performed in triplicate. Student’s t-test, *, P < 0.05; **, P < 0.01; ***, P < 0.001; ****, P < 0.0001. RLU=relative light units.

**Supplementary Figure 4.**
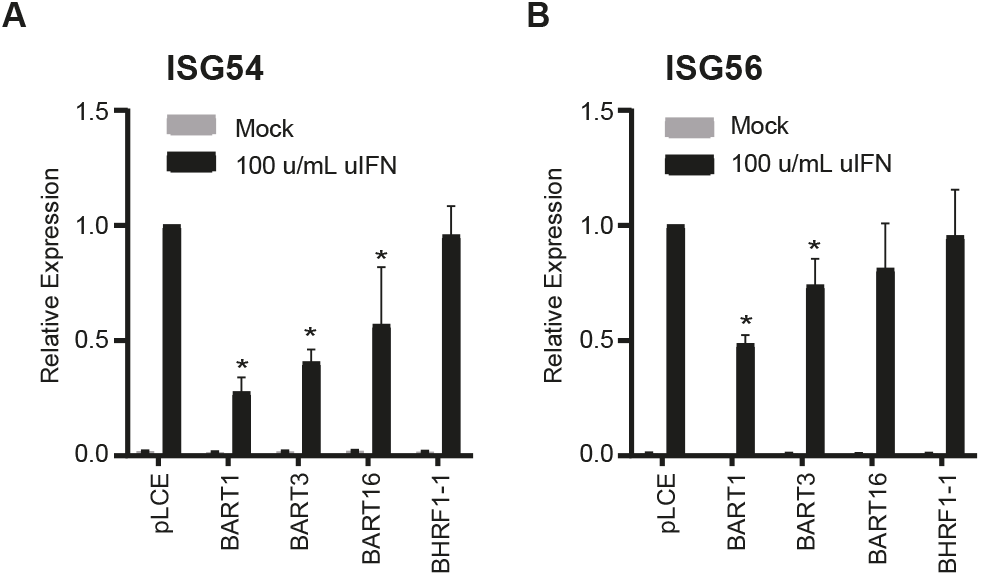
ISG54 and ISG56 are regulated by individual EBV-encoded miRNAs. **(A-B**) HEK293T cells were transfected with either pLCE control or EBV miRNAs. 48 hrs posttransfection, cells were treated with 100 unit/mL uIFN for 6 hrs and total RNA was harvested. ISG54 **(A)** and ISG56 **(B)** expression levels were assayed by qRT-PCR. Values are normalized to GAPDH and shown relative to pLCE control cells treated with uIFN. Reported are the averages of three independent experiments performed in duplicate. Student’s t-test, *p<0.05.

**Supplementary Figure 5.**
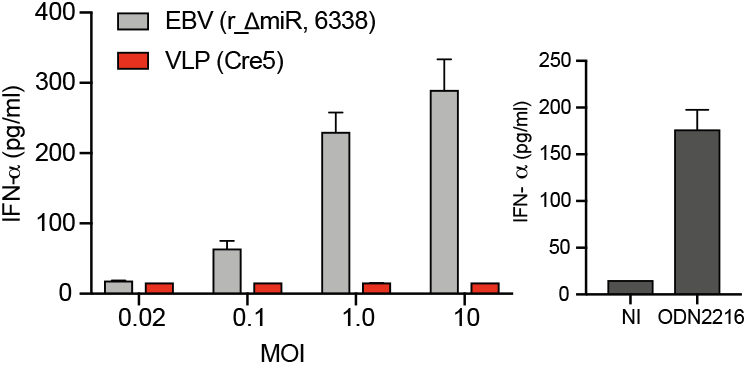
Virus-like particles (VLPs) do not induce IFN-α release. Biological replicate of the experiment shown in Figure 6A. PBMCs were infected with r_ΔmiR EBV or with an equivalent dose of virus-like particles (VLP), which do not contain viral DNA. The culture supernatants were collected 20 h post-infection and IFN-α levels were quantitated by ELISA.

**Supplementary Figure 6.**
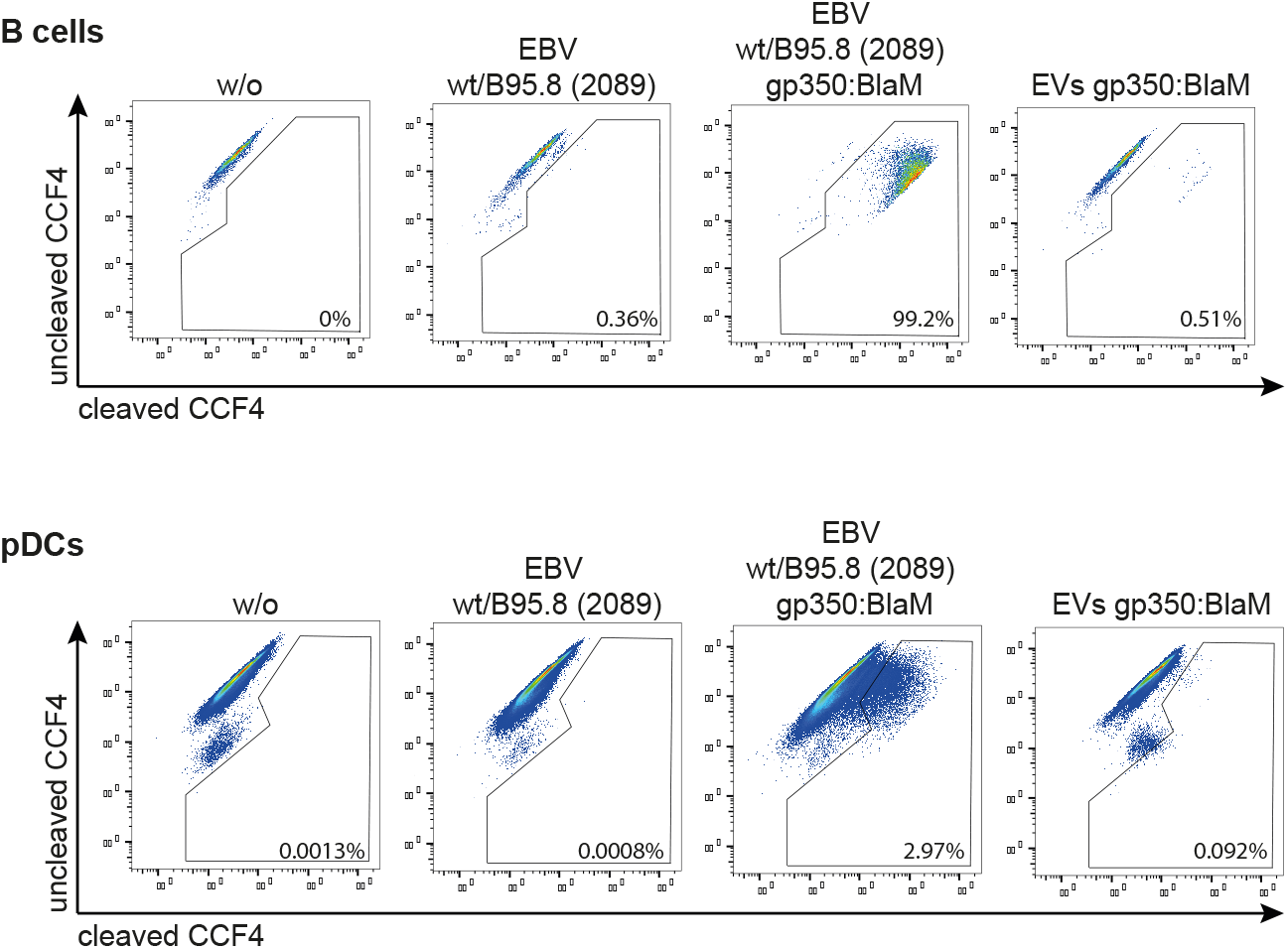
EBV fuses with a minor fraction of pDCs. Representative flow cytometry analysis of B cells and pDCs infected with unmodified wt/B95.8 (2089) EBV or wt/B95.8 (2089) EBV assembled with gp350:BlaM. As negative control cells were left uninfected or were treated with extracellular vesicles (EVs) containing gp350:BlaM. After infection for 4 hours the cells were washed and loaded with CCF4-AM at room temperature for 20 hours. B cells or pDCs were stained with fluorochrome-coupled antibodies directed against CD19 or CD303 and CD304, respectively, and analyzed by flow cytometry.

**Supplementary Figure 7.**
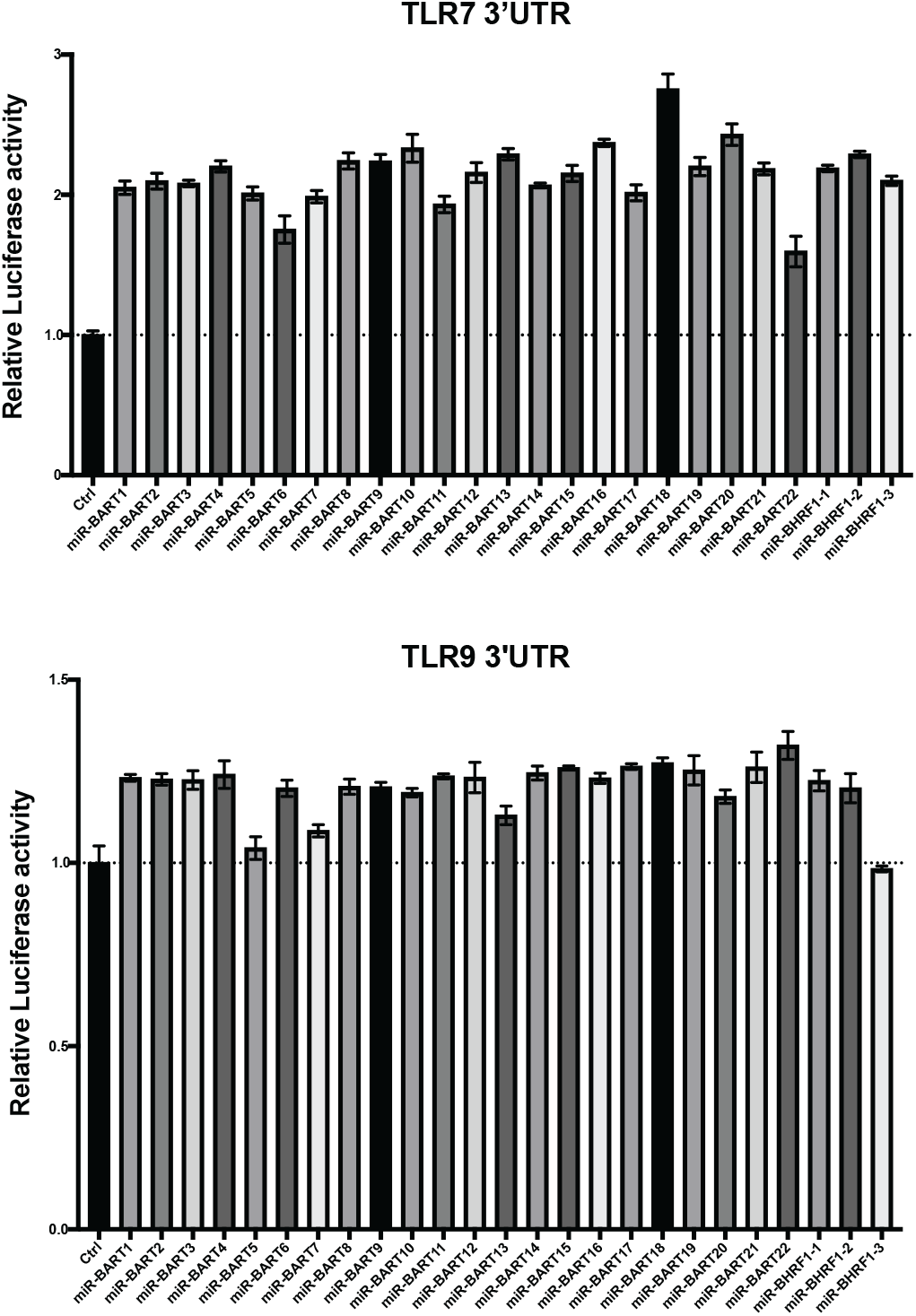
EBV’s miRNAs do not target TLR7 and TLR9 3’UTRs. 293T cells were co-transfected with the indicated wild-type luciferase reporter plasmids (WT) for TLR7 or TLR9 with or without a miRNA-encoding plasmid. Luciferase expression in these cells was assessed and normalized to lysates from cells co-transfected with the wild-type 3’-UTR reporter and the empty miRNA expression plasmid (Ctrl).

