## Supplementary Table 1 for "Multiple viral microRNAs regulate interferon release and signaling early during infection with Epstein-Barr virus"

Overview of EBV strains

| **EBV strain** | **Genotype** | **Specification** | **Genealogy** |
| --- | --- | --- | --- |
| wt/B95.8 (2089) | Wild-type,  13 miRNAs | Reference laboratory EBV strain | B95-8 EBV strain |
| r_wt/B95.8 (6008) | Wild-type,  44 miRNAs | EBV reference strain, deletion in wt/B95.8 (2089) restored | Based on wt/B95.8 (2089) |
| r_ ΔmiR (6338) | Knockout of miRNA coding loci | Scrambled pre-miRNA loci | Based on r_wt/B95.8 (6008) |
| ΔEBER (6431) | Knockout of EBER1 and 2 | Insertional mutagenesis of EBER1,2 locus | Based on r_wt/B95.8 (6008) |
| ΔEBER/ΔmiR (6432) | Knockout of EBER1,2 and miRNA coding loci | Scrambled pre-miRNA loci;  insertional mutagenesis of EBER1,2 locus | Based on r_ ΔmiR (6338) |
| ΔLF2 (6522) | Knockout of LF2 | Stop codon TAA replacing codon 38 in LF2’s open reading frame | Based on r_wt/B95.8 (6008) |
